# Psychopharmacological Effects of Cannabidiol in Rodent Models of Autism Spectrum Disorder: A Systematic Review of Behavioral Outcomes, Neurobiological Mechanisms, and Methodological Quality

**DOI:** 10.64898/2026.10.09.757959

**Authors:** Bruna Rabelo Ribeiro Domingues, Gabriel Santos Félix, Maria Fernanda Lucena Bezerra, Priscila Medeiros, Renato Leonardo de Freitas

**Affiliations:** Laboratory of Neurosciences of Pain & Emotions, Department of Neurosciences and Behavioral Sciences, Ribeirão Preto Medical School, University of São Paulo (FMRP-USP), Ribeirão Preto, SP, Brazil; Graduate Program in Neuroscience, Ribeirão Preto Medical School, University of São Paulo (FMRP-USP), Ribeirão Preto, SP, Brazil; Department of Psychology, Federal University of Paraíba (UFPB), João Pessoa, PB, Brazil; Department of Psychobiology, Ribeirão Preto School of Philosophy, Sciences and Letters, University of São Paulo (FFCLRP-USP), Ribeirão Preto, SP, Brazil; Institute of Natural Sciences, Federal University of Alfenas (UNIFAL-MG), Alfenas, MG, Brazil

**Keywords:** Autism Spectrum Disorder, Cannabidiol, CBD, Endocannabinoid System, Animal Models, Psychopharmacology, Neurodevelopmental Disorders

## Abstract

Autism Spectrum Disorder (ASD) is a neurodevelopmental disorder characterized by persistent deficits in social communication and interaction, as well as restricted and repetitive patterns of behavior. Although increasing evidence suggests that cannabidiol (CBD) may represent a promising therapeutic strategy for ASD, the preclinical literature remains fragmented, with substantial variability in experimental models, treatment protocols, CBD formulations, and reported outcomes. This systematic review (PROSPERO registration: CRD420251177933) aimed to critically evaluate the psychopharmacological effects of CBD in rodent models of ASD, with an emphasis on behavioral outcomes, underlying neurobiological mechanisms, and methodological quality. A systematic search was conducted in PubMed, Embase, Web of Science, Scopus, and ProQuest Dissertations & Theses, with a final sample of 14 articles. Due to the substantial heterogeneity across studies, a qualitative synthesis of results was performed using the Synthesis Without Meta-analysis (SWiM) approach. Overall, CBD and CBD-based preparations produced beneficial effects across multiple ASD-relevant behavioral domains. Enhanced social interaction, reductions in repetitive behaviors, anxiety-like responses, and cognitive impairments were observed in specific models. Neurobiological analyses revealed modulation of the endocannabinoid system, restoration of excitation/inhibition balance, anti-inflammatory and antioxidant effects, and transcriptional regulation. Positive outcomes were observed across pharmacological, genetic, and maternal immune activation models, suggesting that CBD acts on convergent neurobiological pathways implicated in ASD pathophysiology. Despite these promising findings, important limitations were identified, including substantial heterogeneity in experimental protocols and methodological limitations, such as insufficient reporting of randomization, blinding, and allocation concealment procedures, as well as the underrepresentation of female animals. This review synthesizes behavioral, neurobiological, and methodological evidence on CBD and CBD-based preparations in rodent models of ASD, highlighting its therapeutic potential and the need for more rigorous, standardized preclinical investigations that directly compare isolated CBD with its formulations and combinations.

## 1. Introduction

Autism Spectrum Disorder (ASD) is a heterogeneous neurodevelopmental disorder characterized by persistent deficits in social communication and interaction, accompanied by restricted and repetitive patterns of behavior, interests, and activities (American Psychiatric Association, 2022; Mottron, 2020). Although substantial advances have been achieved in understanding the biological basis of ASD, its etiology remains incompletely understood.

Current evidence supports a complex interaction among genetic, epigenetic, and environmental factors contributing to the development and expression of the disorder (Masi et al., 2017; Devlin & Scherer, 2012). Genetic deletions, duplications, and rare variants have been implicated in ASD susceptibility, yet the underlying genetic cause remains unidentified in a considerable proportion of cases. Furthermore, prenatal, perinatal, and postnatal environmental factors have also been associated with ASD risk (Masi et al., 2017).

ASD symptoms typically emerge during early childhood and may become more evident as social demands increase throughout development (Webb & Joness, 2009). Individuals with ASD commonly exhibit impairments in social reciprocity, verbal and non-verbal communication, and the ability to establish and maintain interpersonal relationships. In addition, sensory processing abnormalities, including hyperresponsiveness, hyporesponsiveness, and atypical sensory interests, are frequently reported (Lord et al., 2020; Bal et al., 2019).

Beyond the core diagnostic features, ASD is frequently accompanied by psychiatric and neurological comorbidities, including anxiety disorders, depression, attention-deficit/hyperactivity disorder (ADHD), epilepsy, mood disorders, sleep disturbances, and schizophrenia spectrum disorders (Hossain et al., 2020; Lord et al., 2020). These associated conditions substantially contribute to functional impairment and reduced quality of life, highlighting the need for effective therapeutic strategies capable of addressing both core symptoms and comorbid manifestations.

Despite increasing knowledge regarding ASD pathophysiology, pharmacological treatment options remain limited. Currently available medications primarily target associated symptoms such as irritability and aggression rather than the core deficits in social communication and repetitive behaviors. Moreover, the heterogeneity of ASD and its developmental nature represent significant challenges for the development of effective and broadly applicable interventions (Lord et al., 2020).

In this context, increasing attention has been directed toward the therapeutic potential of cannabinoids. Cannabidiol (CBD), a non-psychoactive phytocannabinoid derived from *Cannabis sativa*, has emerged as a promising therapeutic candidate due to its broad pharmacological profile and favorable safety characteristics (Pereira et al., 2025; Silva et al., 2024). Experimental and clinical studies have demonstrated anxiolytic, anti-inflammatory, antioxidant, anticonvulsant, and neuroprotective properties of CBD, suggesting its potential relevance for neurodevelopmental disorders.

Pharmacologically, the preference for CBD over other cannabinoids, such as Δ9-tetrahydrocannabinol (THC), stems from its unique binding profile and mechanism of action at the CB_1_ receptor. Unlike THC, which acts as an orthosteric agonist, CBD exhibits negligible affinity for this primary binding site. Instead, it functions as a negative allosteric modulator (Thomas et al., 2007; Laprairie et al., 2015). By binding to a secondary allosteric site, CBD attenuates the potency of both endogenous and exogenous agonists without exerting intrinsic activity on the receptor itself. This distinct mechanism is the primary biochemical rationale for CBD’s lack of psychotomimetic effects (Pertwee, 2007).

The endocannabinoid system has been implicated in several biological processes relevant to ASD, including neurodevelopment, synaptic plasticity, emotional regulation, social behavior, and maintenance of excitation/inhibition (E/I) balance within neural circuits (Schnapp et al., 2022; Maldonado et al., 2020). Alterations in endocannabinoid signaling have been reported in both individuals with ASD and animal models of the disorder, supporting the hypothesis that modulation of this system may represent a viable therapeutic strategy.

Consistent with this hypothesis, preclinical studies have demonstrated the beneficial effects of CBD across multiple rodent models of ASD. In BTBR mice, CBD administration reduced repetitive behaviors and improved social interaction (Shrader et al., 2024). Similarly, in valproic acid (VPA)-induced models, CBD prevented the development of autism-like phenotypes and improved social behavior, suggesting both therapeutic and neuroprotective effects (Costa et al., 2024; Taylor et al., 2024). Nevertheless, the available literature remains highly heterogeneous regarding animal models, cannabinoid formulations, doses, routes of administration, treatment duration, and outcome measures (Ornoy et al., 2024; Loss et al., 2021).

While a recent systematic review by Vasconcelos et al. (2026) examined the behavioral and neurological effects of CBD in animal models of ASD, its scope was strictly limited to isolated CBD. Although this approach effectively isolates the compound’s specific effects, it fails to capture the diverse experimental and clinical contexts in which CBD is applied. This limitation is significant, as pharmacological outcomes can vary substantially depending on the intervention. For instance, in non-ASD animal models, co-administering CBD with risperidone, or altering their administration sequence, yielded distinct behavioral, motor, metabolic, and oxidative stress outcomes, highlighting how drug interactions can modulate CBD’s efficacy (Kajero et al., 2022).

Consequently, the primary objective of the present systematic review is to critically synthesize the available evidence regarding the psychopharmacological effects of CBD in rodent models of ASD, focusing on behavioral outcomes, underlying neurobiological mechanisms, and methodological quality. Secondarily, this review aims to evaluate the distinct outcome profiles associated with various CBD interventions, accounting for the diversity of available formulations and compositions.

## 2. Methods

### 2.1 Study Design and Protocol Registration

This systematic review was conducted to evaluate the psychopharmacological effects of cannabidiol (CBD) and CBD-related formulations in rodent models of autism spectrum disorder (ASD). The review followed the Preferred Reporting Items for Systematic Reviews and Meta-Analyses (PRISMA 2020) guidelines.

The review protocol was prospectively registered in the International Prospective Register of Systematic Reviews (PROSPERO) under registration number CRD420251177933. Due to the substantial methodological heterogeneity among the included studies, a quantitative meta-analysis was deemed inappropriate. Consequently, data synthesis was performed in accordance with the Synthesis Without Meta-analysis (SWiM) reporting guideline.

This review was designed to evaluate not only the behavioral effects of CBD in ASD models but also the neurobiological mechanisms underlying these effects, as well as the methodological quality of the available evidence.

### 2.2 Search Strategy

The search strategy was developed according to the PICO framework:

* Population (P): rodent models of autism spectrum disorder;

* Intervention (I): cannabidiol administration, acutely or chronically;

* Comparison (C): vehicle-treated or untreated ASD models;

* Outcomes (O): behavioral (sociability, anxiety, stereotypies, nociceptive changes), and neurobiological effects of CBD.

A systematic search was conducted in PubMed, Embase, Web of Science, Scopus, and ProQuest Dissertations & Theses. Search terms were identified using Medical Subject Headings (MeSH) and adapted according to the indexing requirements of each database. Searches were conducted in October 2025 without restrictions regarding publication year. A PRISMA flowchart was developed in accordance with the methodological guidelines for preparing systematic reviews (PRISMA 2020). The complete search strategy is provided in the Supplementary Material.

### 2.3 Eligibility Criteria

Studies were considered eligible if they:

1. Employed validated rodent models of ASD, including males and females (Panzenhagen et al., 2022);
2. Investigated cannabidiol as the primary intervention;
3. Evaluated behavioral and/or neurobiological outcomes relevant to ASD;
4. Reported original in vivo experimental data;

### Studies were excluded if they

1. Were conducted in vitro or involved human participants;
2. Focused on neurodevelopmental disorders other than ASD;
3. Investigated pharmacological interventions unrelated to CBD;
4. Did not provide sufficient methodological information regarding CBD administration;
5. Were reviews, editorials, conference abstracts, letters, comments, protocols, or non-original publications.

Although the primary focus of this review was cannabidiol (CBD), studies investigating CBD-rich botanical formulations and structurally related phytocannabinoids, such as cannabidivarin (CBDV), were also considered eligible when CBD represented the principal pharmacologically active constituent or when the intervention was specifically developed to investigate CBD-mediated effects.

### 2.4 Study Selection

All records identified through database searches were imported into the Rayyan software for duplicate removal and screening.

The articles were independently analyzed by two reviewers. For the initial title and abstract analysis, exclusion criteria were hierarchically listed, reaching a level of agreement between reviewers of 94%. Disagreements were resolved through the mediation of a third reviewer. After the initial title and abstract analysis, the previously included studies were summarized for full-text analysis. The level of agreement for full-text analysis between reviewers was 98%. Again, a third reviewer was responsible for mediating the disagreements.

Additional eligible articles were identified by analyzing the reference lists of the included studies, using previously defined inclusion and exclusion criteria. A grey literature search was also conducted in the ProQuest Dissertations & Theses database in January 2026, aiming to minimize publication bias and identify relevant studies not indexed in traditional databases, such as theses and dissertations, thereby ensuring a more complete representation of the available evidence on the topic. Eight studies were included for secondary screening, which was performed independently by two reviewers. The agreement level was 99%, and a third reviewer was responsible for mediating any disagreement. The entire process of identifying, selecting, and including these studies is presented in the PRISMA flow diagram (Figure 1).

**Figure 1.**
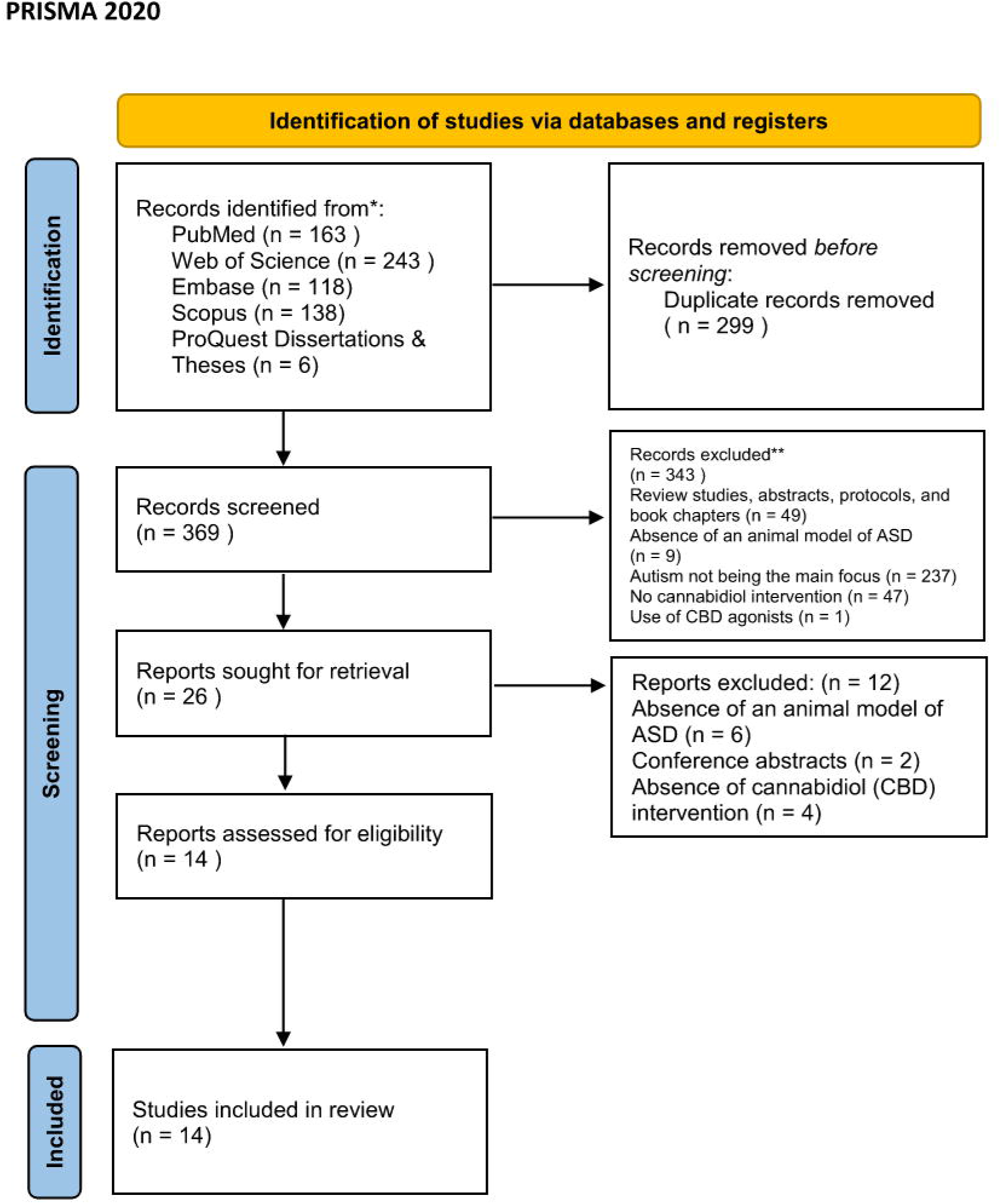
PRISMA flow diagram, with identifying, selecting, and including studies from PubMed, Embase, Web of Science, Scopus, and ProQuest Dissertations & Theses.

### 2.5 Risk of Bias Assessment

Methodological quality was independently evaluated by two reviewers using the SYRCLE Risk of Bias Tool, an adaptation of the Cochrane Risk of Bias framework specifically designed for animal studies (Hooijmans et al., 2014). Each reviewer completed the assessment spreadsheet containing the SYRCLE domains individually. In cases of disagreement among reviewers regarding the classification of bias risk in any domain, a third reviewer acted as a mediator to reach a final consensus.

The results of the risk-of-bias assessment are presented in tabular and graphical formats, highlighting the proportion of studies classified as having low, uncertain, or high risk of bias in each domain. The risk of bias graph was generated using the robvis (Risk of Bias Visualization) tool, developed by McGuinness & Higgins (2021), and is presented as an unweighted bar graph, since this review did not conduct a quantitative meta-analysis. Regarding the judgment, the ‘Overall’ column was added due to a technical requirement of the RobVis tool and populated based on the presence of at least one domain with a high risk of bias (high overall risk) or the predominance of domains with uncertain or low risk (uncertain or low risk, respectively).

The following domains were assessed:

* Sequence generation;

* Baseline characteristics;

* Allocation concealment;

* Random housing;

* Blinding of caregivers and investigators;

* Random outcome assessment;

* Blinding of outcome assessors;

* Incomplete outcome data;

* Selective outcome reporting;

* Other potential sources of bias.

### 2.6 Data Extraction and Synthesis

Data extraction was independently conducted by two reviewers using a standardized extraction form, in accordance with PRISMA 2020 recommendations.

Initially, a standardized data extraction form was developed to ensure that information was collected clearly and consistently across the analyzed studies. Extraction was conducted independently by two reviewers, following the recommendations of Campbell et al. (2020). Discrepancies identified between the reviewers were resolved by a third reviewer with specific expertise in experimental studies and data, who made the final decision. The following information was extracted from each included study:

* Author and year of publication;

* Country of origin;

* Animal species, strain, sex, and age;

* ASD model employed;

* Cannabinoid formulation;

* Dose and route of administration;

* Treatment duration;

* Behavioral outcomes;

* Neurobiological outcomes;

* Principal findings.

Due to substantial heterogeneity across studies in animal models, cannabinoid formulations, dosing regimens, behavioral paradigms, outcome measures, and reporting practices, a quantitative meta-analysis was deemed inappropriate. Therefore, findings were synthesized narratively following SWiM recommendations. In the absence of data or when information was unclear, the corresponding authors of the original studies were contacted for further clarification.

## 3. Results

### 3.1 Study Selection

The findings were synthesized in accordance with SWiM recommendations and organized into six major domains: study characteristics, experimental ASD models, CBD treatment protocols, behavioral outcomes, neurobiological mechanisms, and risk-of-bias assessment.

The database search identified 662 records. After removal of 299 duplicates, 369 studies remained for title and abstract screening. Of these, 343 records were excluded for the following reasons: review studies, abstracts, protocols, and book chapters (49), absence of an animal model of ASD (9), absence of autism as a focus (237), absence of cannabidiol intervention (47), and use of CBD agonists (1). Twenty-six studies underwent full-text assessment, and 14 studies were ultimately included in the qualitative synthesis (Figure 1).

The included studies were published between 2019 and 2025, with a marked increase in publications during 2024 and 2025, reflecting the growing scientific interest in the therapeutic potential of cannabidiol in ASD models.

### 3.2 Characteristics of Included Studies

The 14 included studies were conducted in six distinct countries: Brazil (n = 3; Da Silva et al., 2025; Pedrazzi et al., 2025; Costa et al., 2024), United States (n = 4; Shrader et al., 2024; Ferreira et al., 2023; Staben et al., 2023; Shapiro et al., 2022), Italy (n = 2; Manduca et al., 2024; Zamberletti et al., 2019), Israel (n = 1; Poleg et al., 2021), United Kingdom (n = 1; Loomis et al., 2025), Pakistan (n = 1; Faiza Ali et al., 2025), and Nigeria (n = 1; Udodi et al., 2025), with one additional study conducted in China (Zhang et al., 2023). The observed geographical diversity highlights the international expansion of preclinical research on CBD in ASD. Even so, most studies remain concentrated in developed countries. The data extracted from the eligible studies are summarized in Tables 1 and 2.

**Table 1.**
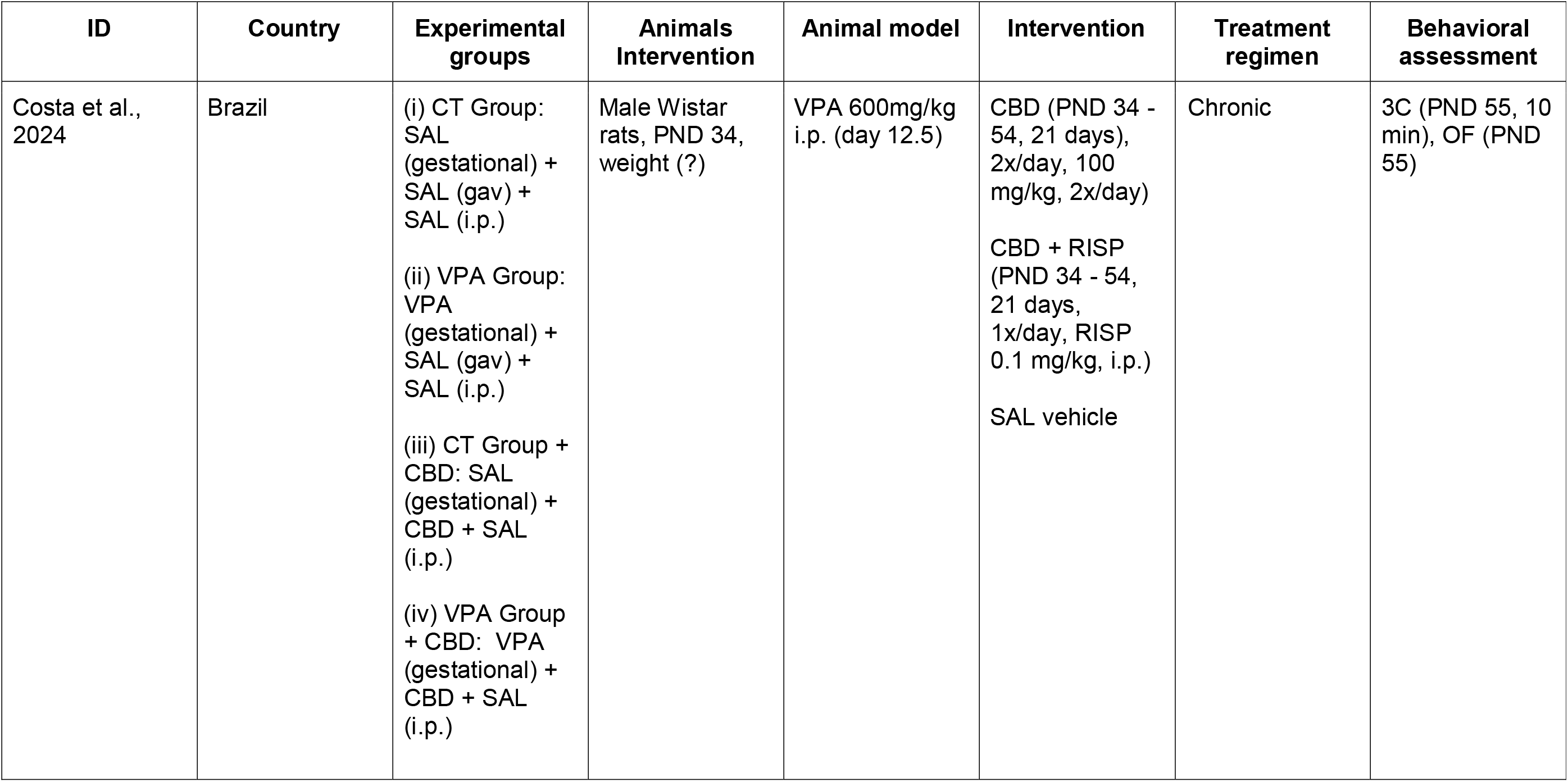
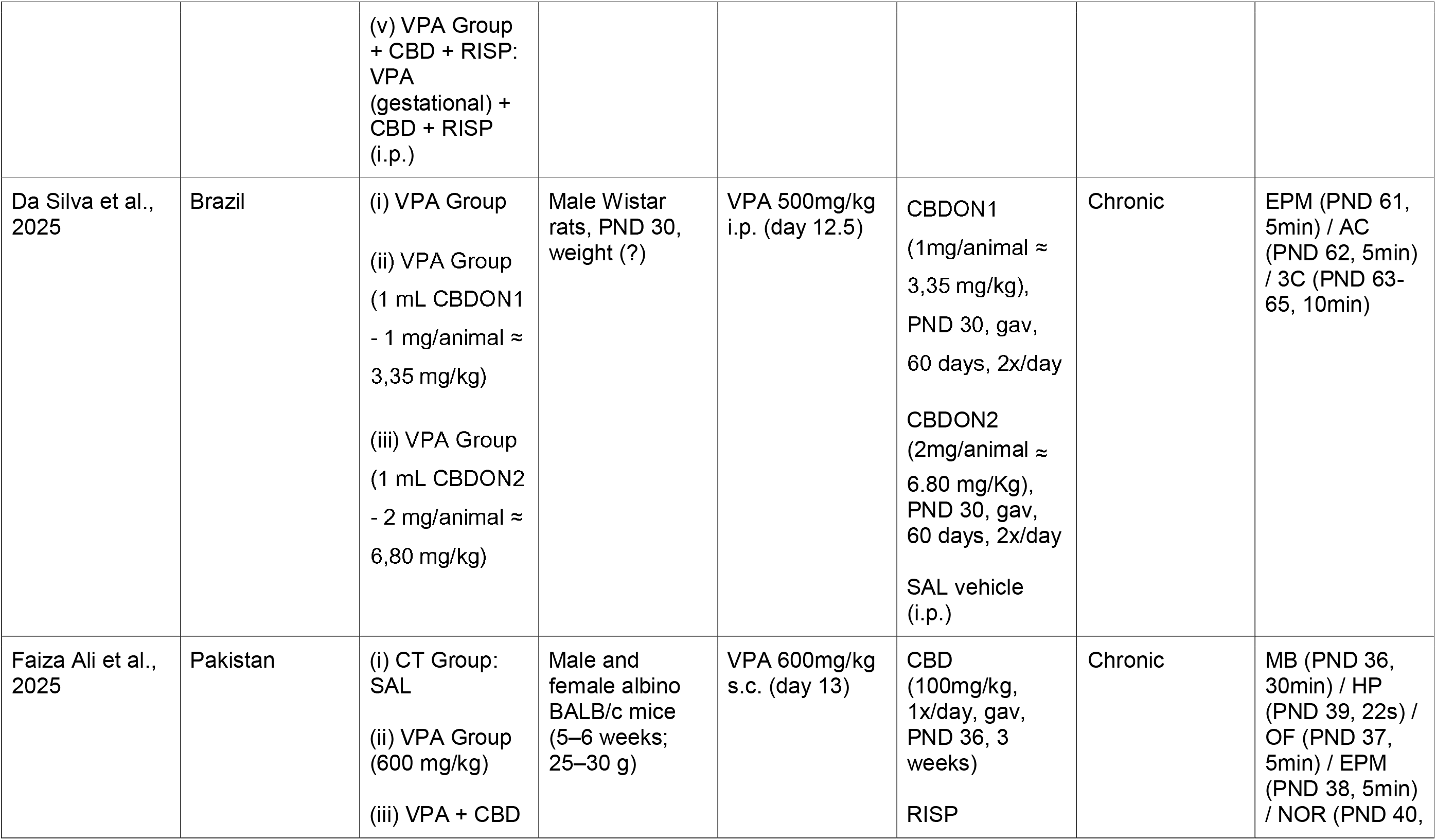
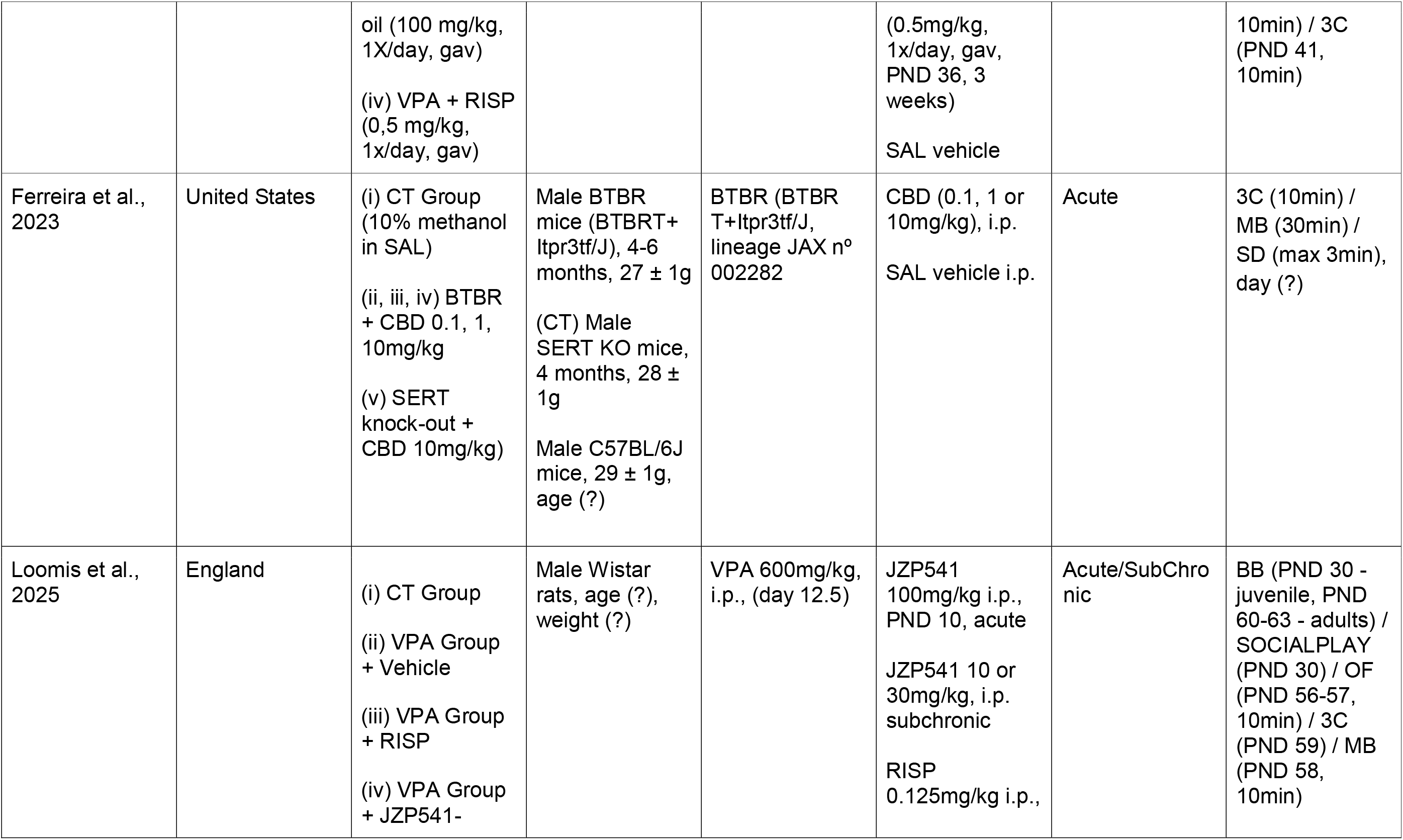
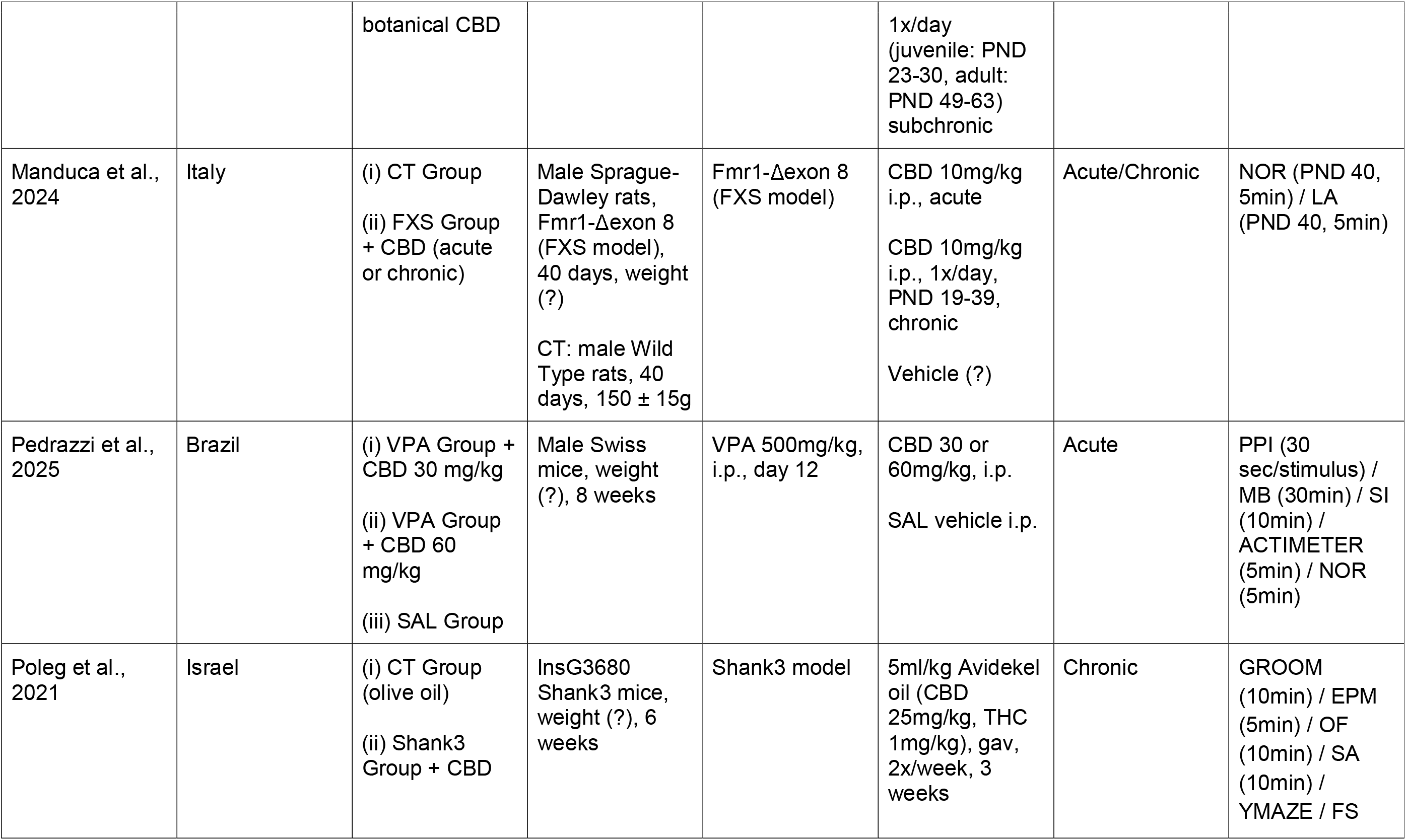
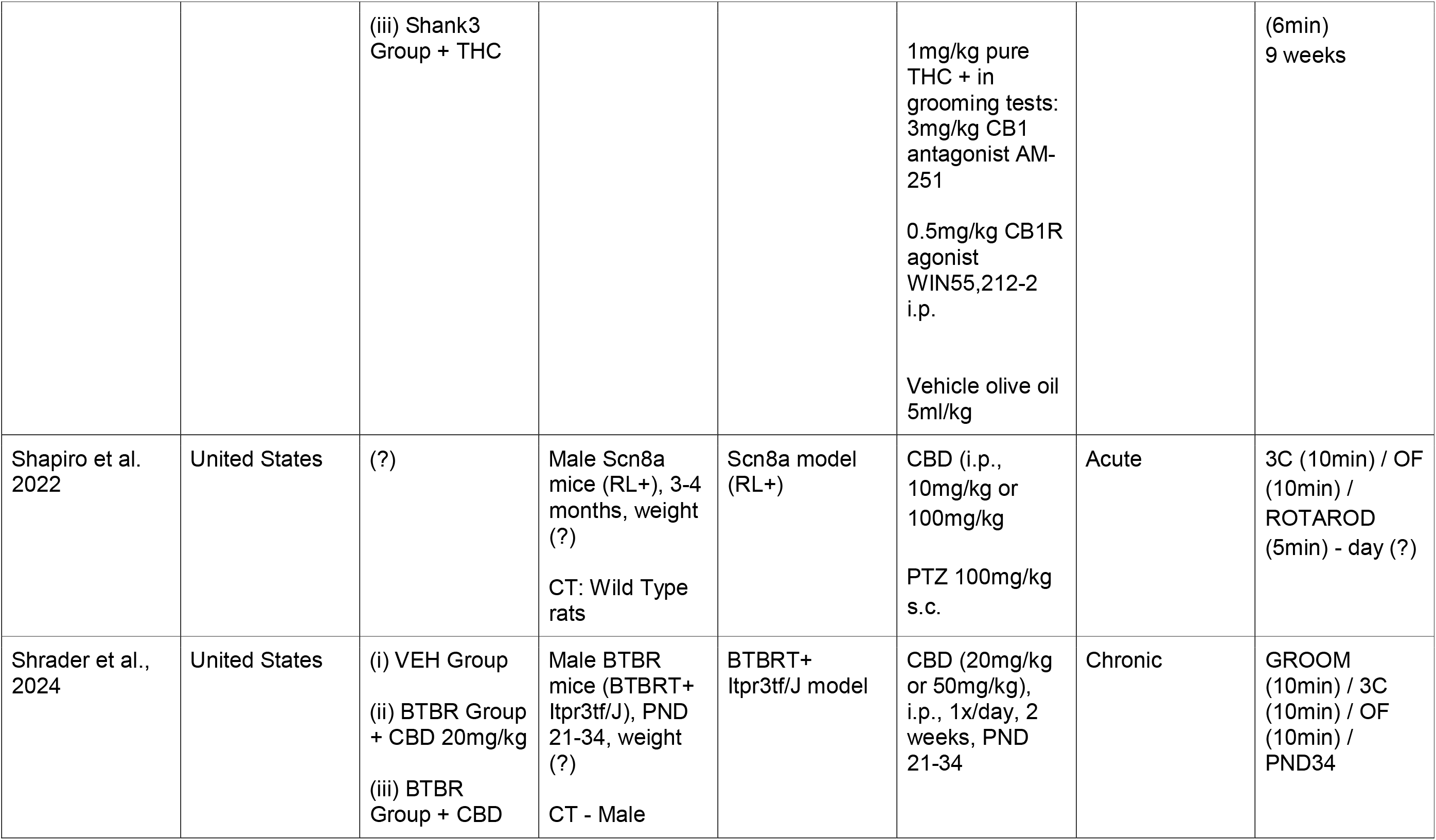
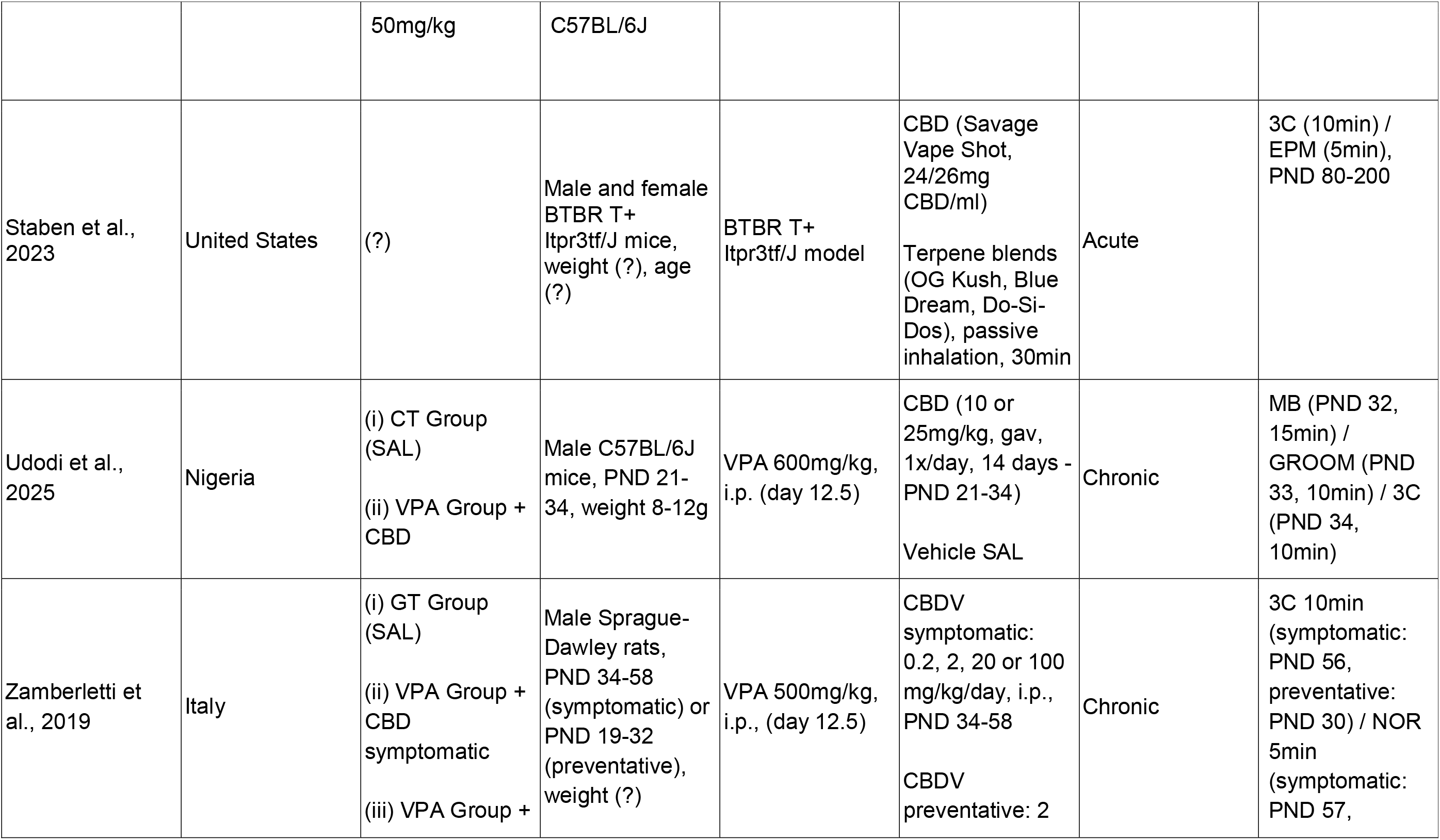
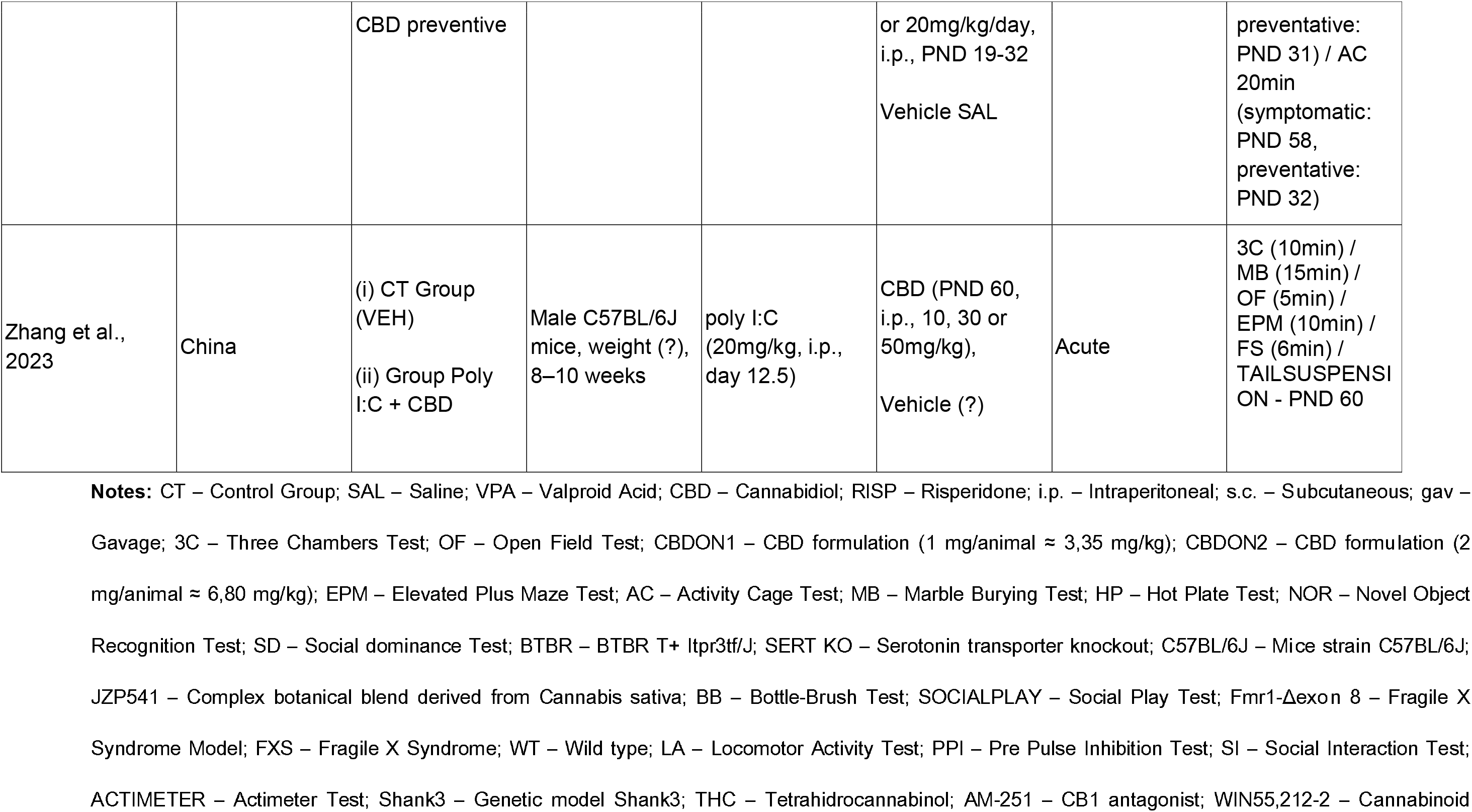
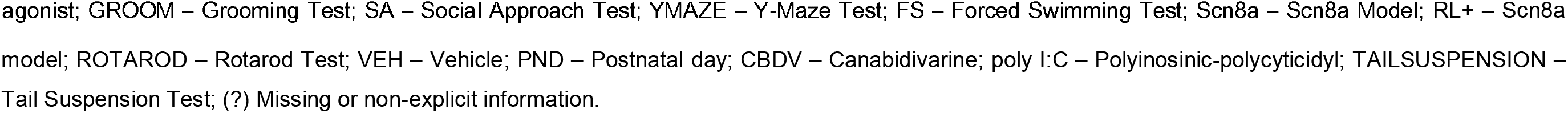
Characterization of the included studies regarding author and year, developing country, experimental groups and animal model implemented, intervention, dosage, and behavioral assessments performed.

**Table 2.**
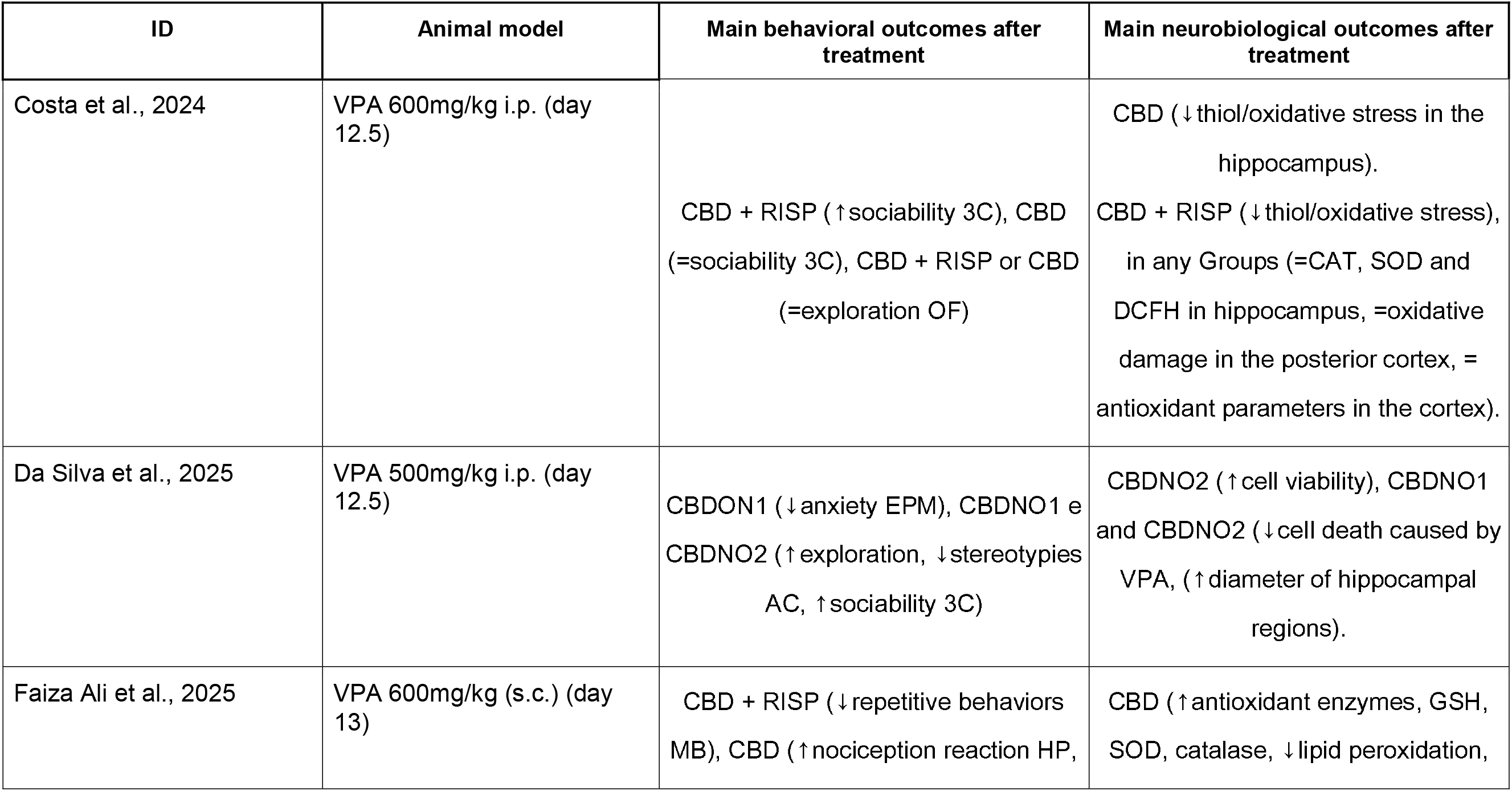
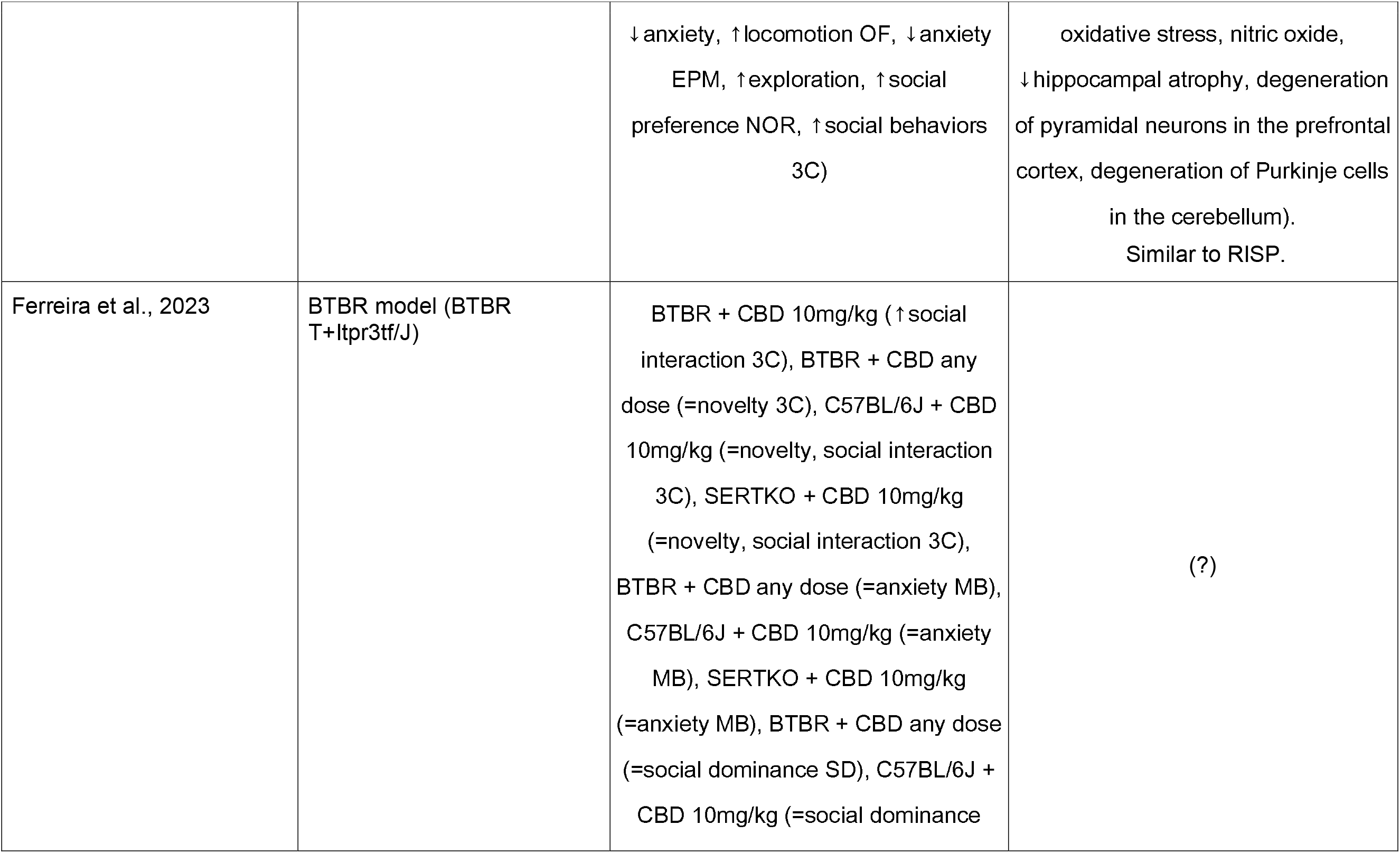
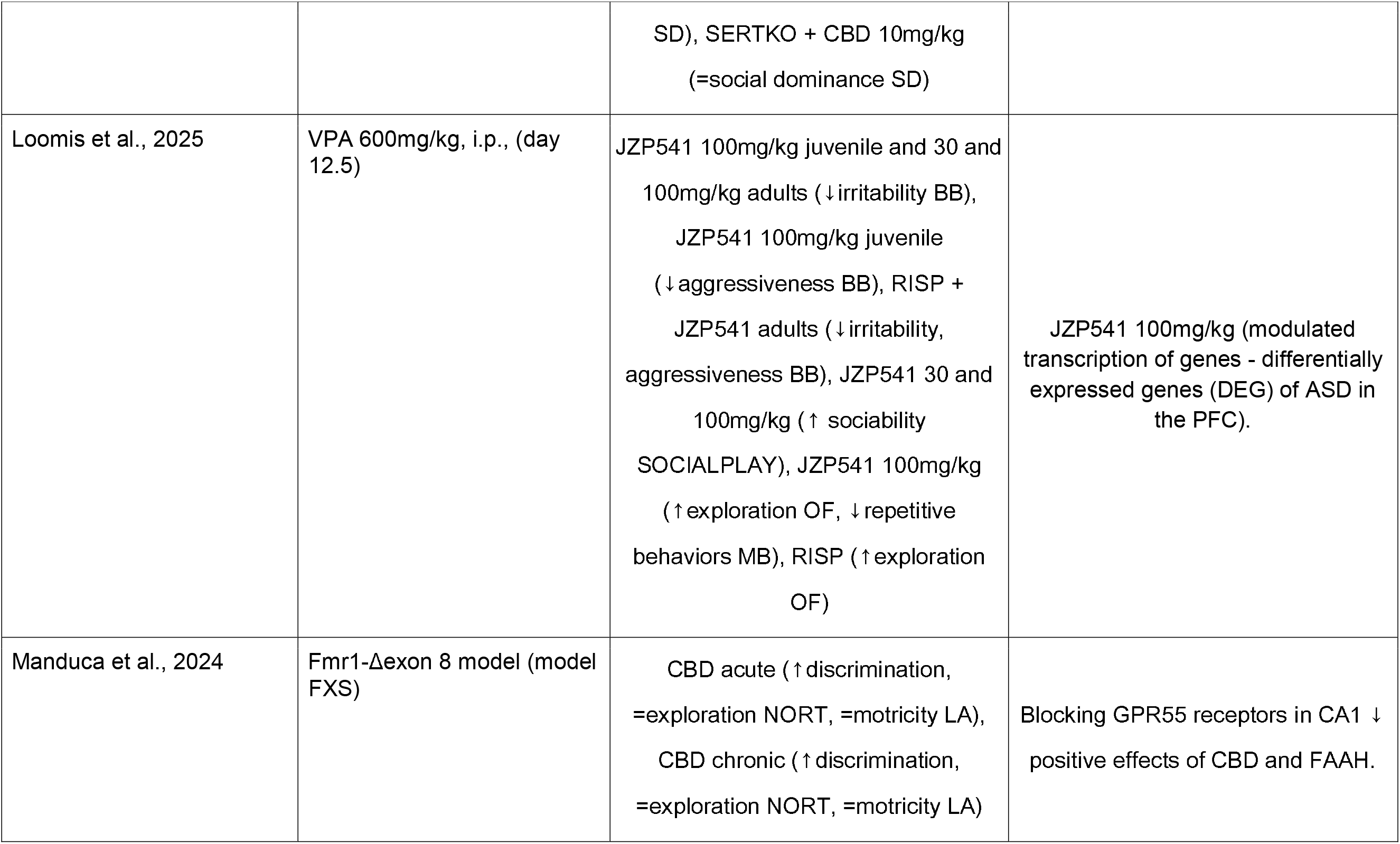
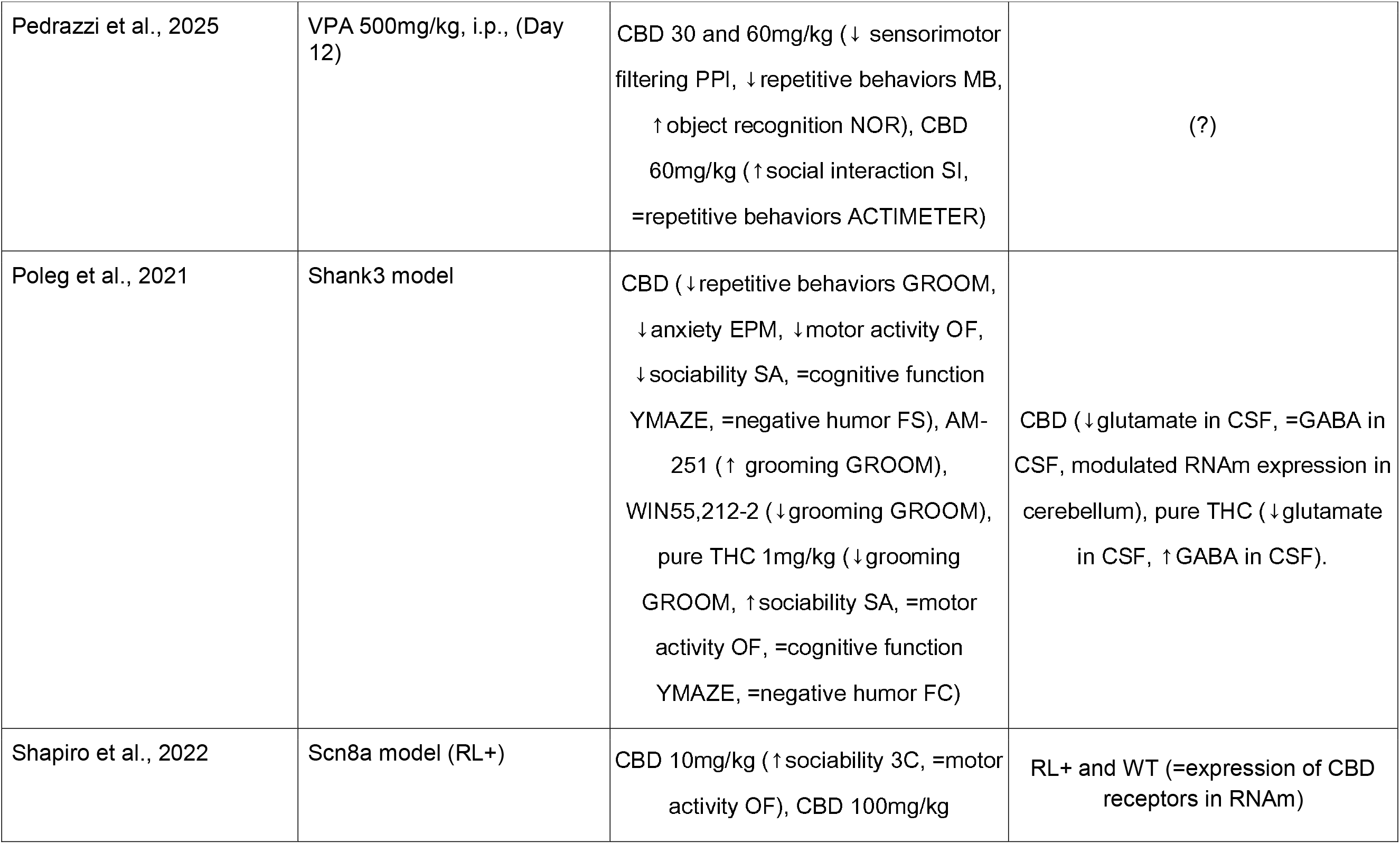
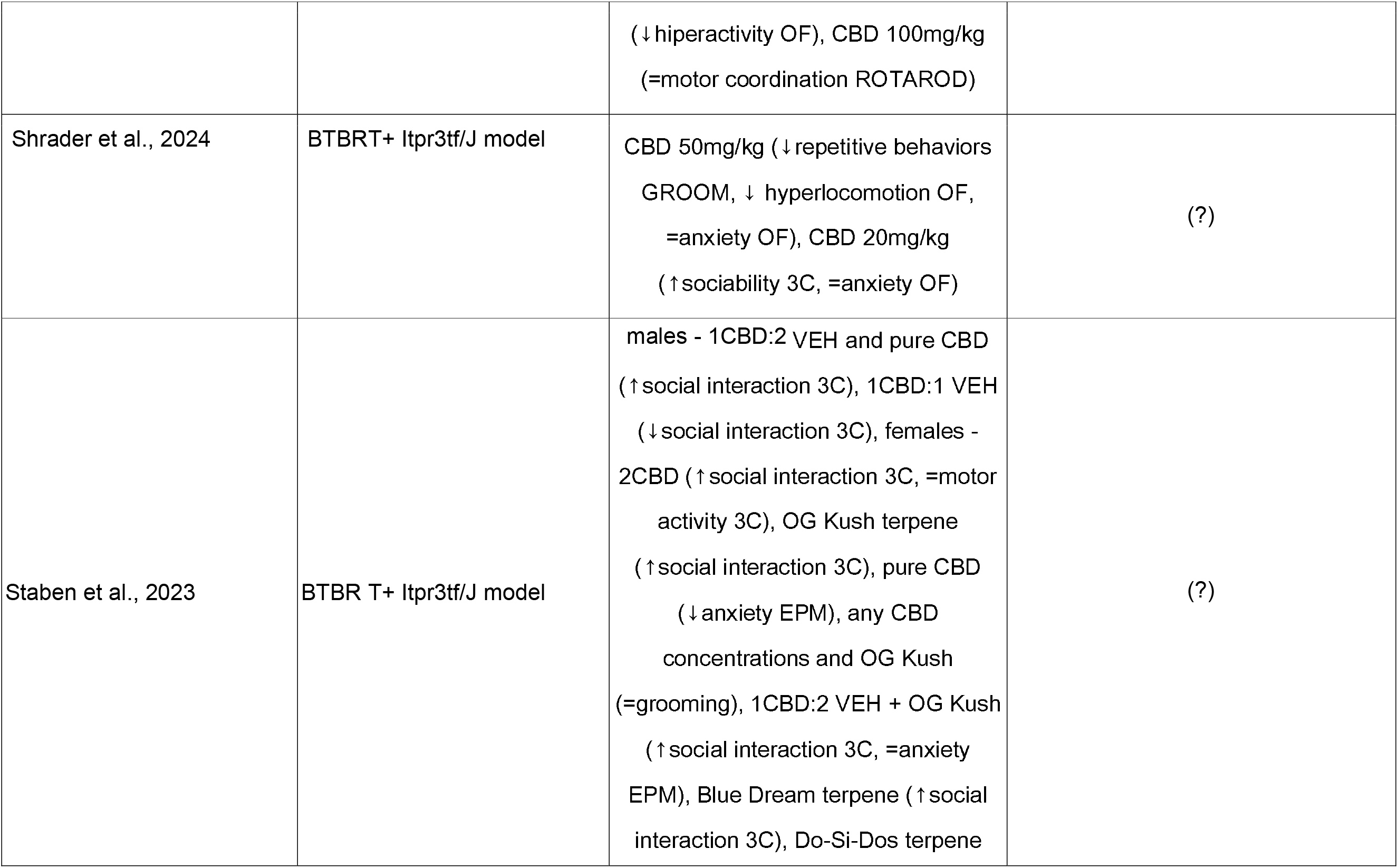
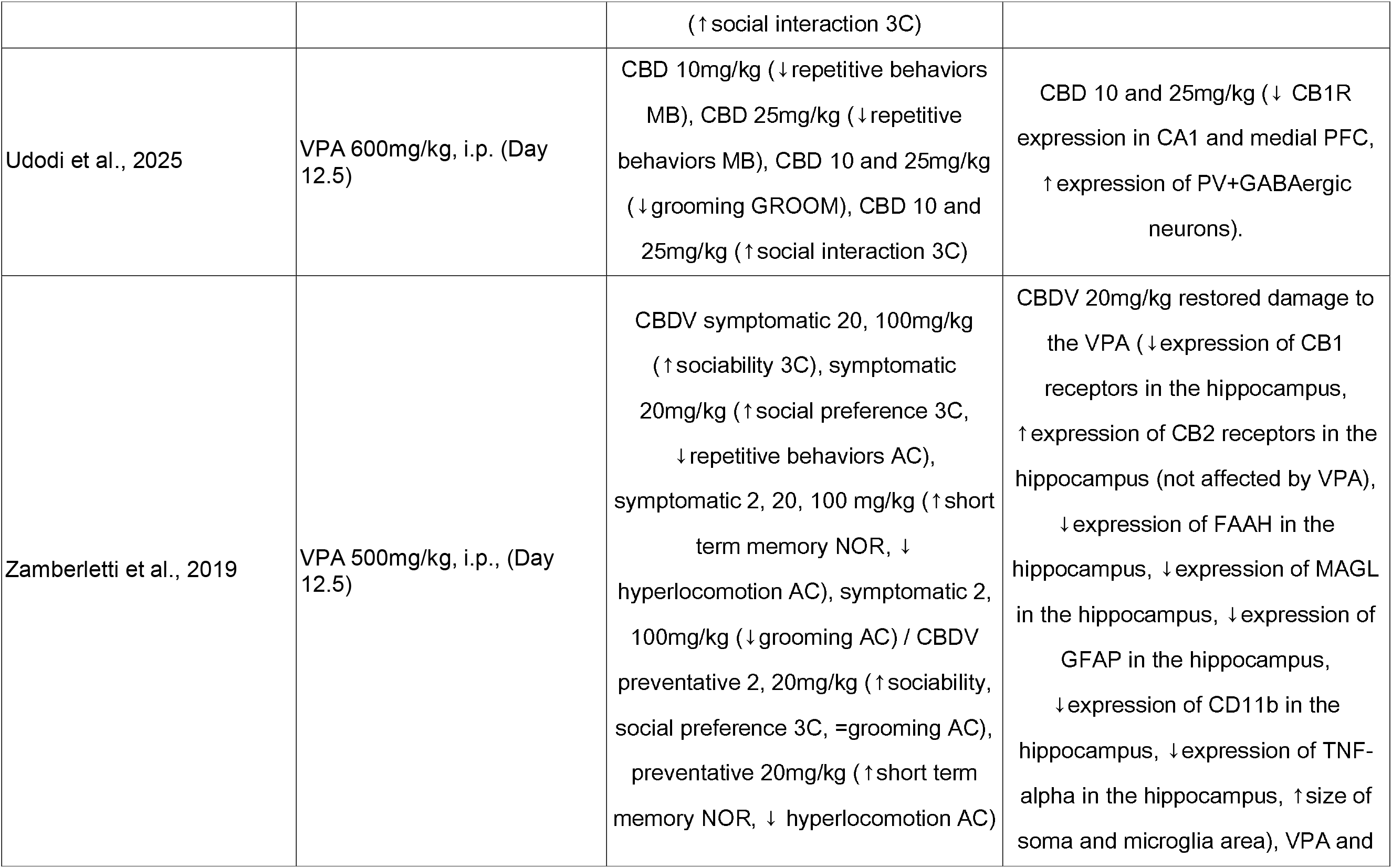
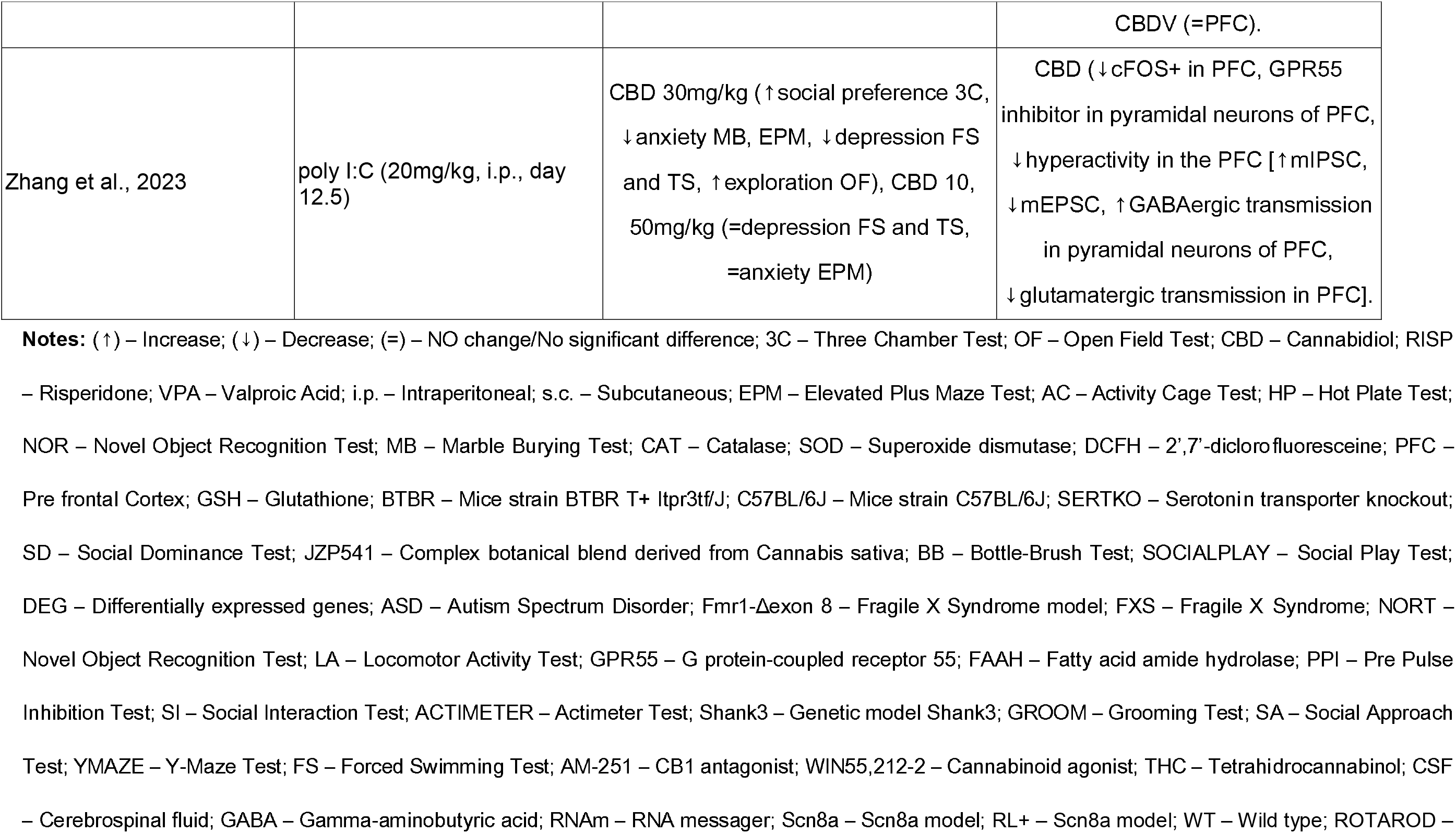
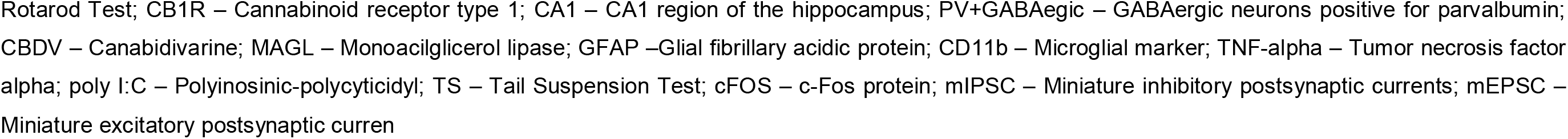
Characterization of the included studies regarding behavioral and neurobiological outcomes after chronic or acute use of CBD and similar substances.

Overall, the included studies demonstrated substantial methodological and experimental heterogeneity regarding ASD models, cannabinoid formulations, dosing regimens, treatment duration, routes of administration, and behavioral assessments. Nevertheless, the consistency of several behavioral and neurobiological outcomes allowed the identification of convergent patterns associated with CBD treatment across distinct experimental paradigms.

### 3.3 Experimental Models of Autism Spectrum Disorder

Three major categories of ASD models were identified among the included studies, reflecting the multifactorial and heterogeneous nature of ASD observed in clinical practice, whose complex neurobiological bases cannot be fully reproduced by a single experimental model.

#### 3.3.1 Valproic Acid Models

Prenatal exposure to valproic acid (VPA) represented the most frequently employed experimental model, used in seven studies included in this review (Da Silva et al., 2025; Faiza Ali et al., 2025; Loomis et al., 2025; Pedrazzi et al., 2025; Udodi et al., 2025; Costa et al., 2024; Zamberletti et al., 2019). Animals exposed to VPA, a widely used antiepileptic and mood-stabilizing drug, displayed behavioral alterations resembling ASD, including deficits in social interaction, repetitive behaviors, anxiety-like responses, sensory abnormalities, and cognitive impairments.

The widespread use of this model reflects its ability to reproduce several behavioral and neurobiological characteristics associated with ASD. However, variability in induction protocols and developmental timing was observed across studies. The doses of VPA used ranged from 500 mg/kg (Da Silva et al., 2025; Pedrazzi et al., 2025; Zamberletti et al., 2019) to 600 mg/kg (Faiza Ali et al., 2025; Loomis et al., 2025; Udodi et al., 2025; Costa et al., 2024), administered intraperitoneally (i.p.) or subcutaneously (s.c.) on gestational days 12, 12.5, or 13. Although the VPA model is widely validated and extensively used in the literature, it has important limitations: (1) it predominantly reproduces the phenotype of syndromic autism associated with teratogenic exposure; (2) most of the included studies used only male rodents, which limits the generalizability of the results considering the existence of ASD in both sexes; and (3) variability in induction protocols between laboratories, in terms of dose and day of application, which makes direct comparison of results difficult.

#### 3.3.2 Genetic Models

Several investigations employed genetic models, including BTBR mice, Shank3-mutant mice, Scn8a-mutant mice, serotonin transporter (SERT) knockout mice, and Fragile X syndrome (Fmr1) models.

The BTBR T+Itpr3tf/J (BTBR) mice were used in three studies (Shrader et al., 2024; Ferreira et al., 2023; Staben et al., 2023). It is an innate lineage that exhibits a robust autism phenotype, including marked reduction in social behavior, increased repetitive and stereotyped behaviors, such as excessive grooming, and learning deficits. BTBR mice exhibit absence of the corpus callosum, reduction of the hippocampal commissure, hyperserotonemia, and alterations in the endocannabinoid system (Meyza & Blanchard, 2017).

Ferreira and colleagues (2023) additionally used SERT KO (serotonin transporter knockout) mice, which exhibit increased synaptic serotonin and sociability deficits. Manduca et al. (2024) used the Fmr1-Δexon8 model (mice with a deletion of exon 8 of the Fmr1 gene), which mimics Fragile X syndrome (FXS), the most common monogenic cause of intellectual disability and the genetic syndrome most frequently associated with ASD. Although the association between FXS and ASD varies across clinical studies, this model remains highly relevant from a translational perspective because it exhibits behavioral phenotypes and characteristics frequently observed in individuals with ASD, including social deficits, stereotypies, sensory alterations, and cognitive deficits. Thus, even in the face of inconsistencies in the literature, Fmr1-based models remain valuable for investigating neurobiological and behavioral mechanisms related to ASD.

In this context of genetic models, Poleg et al. (2021) used InsG3680 Shank3 mice, carriers of a gain-of-function mutation in the SHANK3 gene, which encodes an essential protein of postsynaptic densities of excitatory synapses and is commonly implicated in syndromic autism. Shapiro et al. (2022) used the Scn8a (RL+) model, representing a loss-of-function mutation in the Nav1.6 sodium channel, which is associated with epilepsy and certain characteristics of ASD.

#### 3.3.3 Maternal Immune Activation Models

One study employed maternal immune activation induced by polyinosinic-polycytidylic acid (poly I: C), a model designed to mimic prenatal inflammatory exposure (Zhang et al., 2023). Animals were exposed to maternal immune activation on gestational day 12.5, at a 20mg/kg i.p. dose, and their adult offspring exhibited social deficits, anxiety-like behaviors, and alterations in cortical excitatory and inhibitory neurotransmission, consistent with deficits seen in ASD symptomatology.

### 3.4 Cannabinoid Doses and Administration Routes

Considerable variability was observed regarding cannabinoid doses and treatment protocols. Ranges from 0.1 mg/kg to 100 mg/kg were identified, reflecting the lack of dosage standardization in the preclinical protocols. The main routes of administration were: intraperitoneal (i.p.) (Pedrazzi et al., 2025; Manduca et al., 2024; Shrader et al., 2024; Ferreira et al., 2023; Zhang et al., 2023; Shapiro et al., 2022; Zamberletti et al., 2019), gavage(gav)/oral route (Da Silva et al., 2025; Faiza Ali et al., 2025; Udodi et al., 2025; Costa et al., 2024; Poleg et al., 2021) and passive inhalation (Staben et al., 2023). Loomis et al. (2025) used i.p. with a botanical CBD formulation (JZP541).

Regarding the administration protocol, eight studies used chronic treatment (ranging from 14 days to approximately 9 weeks), four employed acute administration (single dose), and two used both protocols (Manduca et al., 2024; Loomis et al., 2025). It is noteworthy that a direct comparison between acute and chronic administration, when conducted within the same study, yields more robust evidence regarding the need for continuous treatment to achieve lasting effects.

The marked heterogeneity in treatment regimens highlights the absence of standardized protocols for cannabinoid administration in preclinical ASD research.

#### 3.4.1 Acute versus Chronic Administration

Studies with acute administration have demonstrated significant behavioral effects even after a single dose, suggesting rapid-onset mechanisms of action. Ferreira et al. (2023) observed that CBD 10 mg/kg (i.p., acute) increased social interaction in BTBR mice in the three-chamber test (3C), although lower doses (0.1 and 1 mg/kg) did not produce similar effects in this test. At relatively intermediate doses, Zhang et al. (2023) identified a therapeutic window for CBD 30 mg/kg (acute), which reduced repetitive behaviors, anxiety, and depressive-like behaviors in poly I: C mice, while doses of 10 or 50 mg/kg were unable to replicate these effects. Pedrazzi et al. (2025) demonstrated that doses of 30 and 60 mg/kg (acute) reduced repetitive behaviors and improved object recognition. In addition, improvement in social interaction was observed with the higher dose of CBD (60 mg/kg).

For the chronic application protocol, studies have shown variable response profiles depending on the dose, duration, species, strain, and experimental model used. Costa et al. (2024) observed that CBD 100 mg/kg (gav, for 21 days) did not improve sociability in the VPA model on its own. Still, the combination of CBD with risperidone (RISP) produced a significant improvement, suggesting pharmacological potentiation. Udodi et al. (2025) demonstrated that CBD 10 and 25 mg/kg (gav, for 14 days) reduced repetitive behaviors and improved sociability in VPA mice. Zamberletti et al. (2019) obtained significant results with CBDV (a CBD analog) at doses of 2, 20, and 100 mg/kg (i.p., chronic, symptomatic regimen), with the 20 mg/kg dose demonstrating the best efficacy.

Overall, the available evidence indicates that both acute and chronic CBD administration can produce beneficial effects in ASD models. However, chronic treatment appears more consistently associated with widespread behavioral and molecular improvements.

#### 3.4.2 Special Formulations: Nanoemulsions, Botanical Extracts, and Inhalation

In addition to purified CBD, several studies investigated alternative cannabinoid formulations designed to improve bioavailability or explore potential synergistic interactions among phytocannabinoids and terpenes.

Da Silva et al. (2025) investigated the use of CBD nanoemulsion (CBDON) in enriched, at doses of 1 and 2 mg (equivalent to very low doses compared to the others). Both formulations reduced VPA-induced hippocampal cell death and improved behavioral parameters, with CBDON2 (2 mg) exhibiting greater efficacy. The use of nanoemulsion represents a relevant pharmaceutical innovation, with the potential to increase the oral bioavailability of CBD. This benefit is especially important because CBD has low aqueous solubility and undergoes extensive first-pass metabolism, both of which limit its absorption.

Staben et al. (2023) were the only ones to use passive inhalation of CBD with and without terpene mixtures (OG Kush, Blue Dream, Do-Si-Dos). In general, the study showed that CBD increased prosocial behaviors in the animal model used. Furthermore, terpenes mixtures also promoted pro-social effects, both in isolation and in combination with CBD, resulting in more robust effects. The authors also reported that these benefits depend on the combination of different terpenes present in the formulations and occur independently of anxiety-related changes.

Finally, Poleg et al. (2021) used Avidekel, a botanical extract with a CBD: THC ratio of approximately 25:1, administered by gavage for 3 weeks. Although the effects were predominantly attributed to CBD based on CB1 receptor antagonism experiments, the presence of THC and other phytocannabinoids prevents the exclusive attribution of the effects to isolated CBD. Loomis et al. (2025) used JZP541, a botanical formulation of CBD, which demonstrated good results in the VPA model, reinforcing the effectiveness of non-synthetic preparations.

Collectively, these studies suggest that formulation characteristics may significantly influence the pharmacological profile of CBD and represent an important consideration for future translational and clinical investigations.

### 3.5 Behavioral Outcomes

The studies included in this review employed a wide variety of behavioral tests to assess the core and associated domains of ASD in rodents, highlighting deficits in sociability, repetitive and stereotyped behaviors, as well as associated symptoms such as anxiety and memory impairments.

#### 3.5.1 Social Behavior

Improvement in social behavior was the most consistently reported effect across studies.

The Three-Chamber Social Test (3C) test was the most frequently used behavioral instrument, present in 11 of the 14 studies analyzed (Da Silva et al., 2025; Faiza Ali et al., 2025; Loomis et al., 2025; Udodi et al., 2025; Costa et al., 2024; Shrader et al., 2024; Ferreira et al., 2023; Staben et al., 2023; Zhang et al., 2023; Shapiro et al., 2022; Zamberletti et al., 2019). This test is widely recognized in the preclinical literature for assessing sociability and social preference in rodents, allowing measurement of both the approach to the social stimulus and the ability to discriminate between familiar and novel individuals.

Other paradigms, such as the Social Play (Loomis et al., 2025), Social Approach (Poleg et al., 2021), Social Interaction (Pedrazzi et al., 2025), and Social Dominance (Ferreira et al., 2023), were used in one study each.

CBD has demonstrated a prosocial effects in VPA models (Da Silva et al., 2025; Faiza Ali et al., 2025; Loomis et al., 2025; Udodi et al., 2025; Zamberletti et al., 2019), BTBR (Shrader et al., 2024; Ferreira et al., 2023; Staben et al., 2023), Scn8a RL+ (Shapiro et al., 2022) and poly I:C (Zhang et al., 2023). On the other hand, Costa et al. (2024) did not observe an improvement in sociability with CBD alone in the VPA model in Wistar rats; the effect occurred only with the combination of CBD and risperidone. Poleg et al. (2021) reported a reduction in sociability in the Shank3 model with Avidekel, an effect that may be attributed to the THC present in the formulation or the dose used.

The discrepancies found in the study results regarding sociability can be explained by differences in experimental models, doses, routes of administration, species used, test protocols, and, critically, by the non-linear dose-response relationship (inverted U shape), characteristic of CBD, whereby intermediate doses are more effective than very low or very high doses.

#### 3.5.2 Repetitive and Stereotyped Behaviors

Several studies reported reductions in repetitive and stereotyped behaviors following CBD treatment.

At an intermediate frequency, the grooming test was evaluated in 5 studies (Udodi et al., 2025; Shrader et al., 2024; Staben et al., 2023; Poleg et al., 2021; Zamberletti et al., 2019) as a measure of repetitive behaviors.

The Marble Burying (MB) test was used in 7 studies (Faiza Ali et al., 2025; Loomis et al., 2025; Pedrazzi et al., 2025; Udodi et al., 2025; Shrader et al., 2024; Ferreira et al., 2023; Zhang et al., 2023) and is widely adopted as a measure of repetitive and compulsive behaviors, one of the core domains of ASD.

The Activity Cage (AC) was used by Da Silva et al. (2025) and Zamberletti et al. (2019) to assess locomotor activity and stereotyped behaviors.

CBD was effective in reducing repetitive behaviors in several models: VPA (Faiza Ali et al., 2025; Loomis et al., 2025; Pedrazzi et al., 2025; Udodi et al., 2025; Zamberletti et al., 2019), BTBR (Shrader et al., 2024; Staben et al., 2023 [with no effect of CBD on grooming]), and poly I: C (Zhang et al., 2023). Poleg et al. (2021) observed a reduction in grooming in the Shank3 model with Avidekel; however, this effect was partially mediated by the CB1 pathway, as evidenced by increased grooming with the antagonist AM-251 and reduced grooming with the agonist WIN55,212-2.

#### 3.5.3 Anxiety-Like Behaviors

Anxiolytic effects were observed in multiple studies employing open field and elevated plus maze paradigms.

The Open Field (OF) test was the second most used, present in 9 studies (Faiza Ali et al., 2025; Loomis et al., 2025; Costa et al., 2024; Shrader et al., 2024; Ferreira et al., 2023; Staben et al., 2023; Zhang et al., 2023; Shapiro et al., 2022; Poleg et al., 2021), mainly used to assess locomotor and exploratory activity, in addition to providing an indirect index of anxiety behavior.

The Elevated Plus Maze (EPM) was present in 5 studies (Da Silva et al., 2025; Faiza Ali et al., 2025; Staben et al., 2023; Zhang et al., 2023; Poleg et al., 2021) as a classic instrument for assessing anxiety.

Faiza Ali et al. (2025) observed a reduction in anxiety in EPM and OF; Zhang et al. (2023) identified an anxiolytic effect with a dose of 30 mg/kg, but not with 10 or 50 mg/kg; Poleg et al. (2021) reported a reduction in anxiety in the Shank3 model; Pedrazzi et al. (2025) verified a reduction in sensorimotor gating (using the Pre Pulse Inhibition Test, PPI) as an indicator of anxiety; Staben et al. (2023) demonstrated an anxiolytic effect in EPM with CBD by inhalation.

#### 3.5.4 Cognitive Performance

A subset of studies investigated cognitive outcomes. The Novel Object Recognition Test (NOR) was used in 4 studies (Faiza Ali et al., 2025; Pedrazzi et al., 2025; Manduca et al., 2024; Zamberletti et al., 2019) to assess short-term recognition memory.

The Y-Maze Test was also used in one study (Poleg et al., 2021), aiming to assess short-term memory and exploratory behavior.

Manduca et al. (2024) demonstrated improved object discrimination with both acute and chronic administration of CBD 10 mg/kg in Fmr1 KO rats (FXS model), without alterations in locomotor activity. Zamberletti et al. (2019) observed improved short-term memory with CBDV at multiple doses in the symptomatic regimen. Similarly, Pedrazzi et al. (2025) reported improved object recognition with CBD 30 and 60 mg/kg (acute) in VPA Swiss mice.

#### 3.5.5 Locomotor Activity

Shapiro et al. (2022) used the Rotarod paradigm to assess motor function and coordination. The Activity Cage (AC) test, as mentioned, was also intended to evaluate motor activity in mice (Da Silva et al., 2025; Zamberletti et al., 2019).

In general, the doses of CBD used did not significantly affect spontaneous motor activity, except for Poleg et al. (2021), who observed a reduction in motor activity with Avidekel in the Shank3 model, and Shapiro et al. (2022), who found a reduction in hyperactivity with CBD 100 mg/kg in Scn8a mice. Shrader et al. (2024) found that CBD 50 mg/kg (but not 20 mg/kg) reduced hyperlocomotion in BTBR. This observation is particularly important because it indicates that improvements in social and anxiety-related behaviors were unlikely to result from sedation or generalized motor suppression.

#### 3.5.6 Other behaviors

Several studies also used other behavioral tests, focused on specific outcomes. Zhang et al. (2023) and Poleg et al. (2021) used the Forced Swimming Test (FS) as an indicator of depressive-like behaviors. Zhang et al. (2023) also used the Tail Suspension Test as another paradigm for screening depressive symptoms.

Loomis et al. (2025) used the Bottle-Brush Test to evaluate aggressive and irritable behaviors. Tactile perception was also studied using the Prepulse Inhibition Test (PPI) (Pedrazzi et al., 2025) and the Hot Plate Test (Faiza Ali et al., 2025), which evaluate sensory information processing by exposing participants to a loud noise or a hot surface.

### 3.6 Neurobiological Outcomes

Seven of the 14 included studies investigated neurobiological mechanisms underlying the effects of CBD, encompassing the endocannabinoid, glutamatergic, GABAergic, inflammatory, oxidative, and epigenetic systems.

#### 3.6.1 Modulation of the Endocannabinoid System

Zamberletti et al. (2019) demonstrated that CBDV restored alterations in the hippocampal endocannabinoid system induced by VPA by normalizing the reduced expression of the CB1 receptor, increasing the expression of the CB2 receptor (unaffected by VPA), and restoring the expression of the endocannabinoid degradation enzymes FAAH and MAGL in the hippocampus. These effects were accompanied by a reduction in neuroinflammatory markers (GFAP, CD11b, TNF-α) and restoration of microglial morphology, suggesting that hippocampal endocannabinoid modulation is a central mechanism of CBDV’s effects in this model.

Manduca et al. (2024) demonstrated, through selective pharmacological blockade, that the procognitive effects of CBD in the Fmr1 KO model are mediated by the GPR55 receptor in CA_1_ neurons of the hippocampus, with participation of the FAAH pathway. Blockade of hippocampal GPR55 abolished the beneficial effects of CBD on object discrimination, indicating that this receptor, whose involvement in ASD is still poorly explored, may represent a relevant therapeutic target.

Udodi et al. (2025) demonstrated that CBD 10 and 25 mg/kg (chronic) reduced the expression of the CB_1_ receptor in the CA_1_ regions of the hippocampus and in the medial prefrontal cortex of VPA mice. They increased the expression of parvalbumin-positive (PV+) GABAergic interneurons, suggesting restoration of excitation/inhibition (E/I) balance as a mechanism of action. The deficit of PV+ interneurons is one of the most replicated neurobiological findings in both animal and human models of ASD.

#### 3.6.2 Restoration of Excitation/Inhibition Balance

Studies also reported neurobiological effects consistent with restoration of E/I balance.

Zhang et al. (2023) provided direct electrophysiological evidence that CBD modulates the E/I balance in the prefrontal cortex (PFC): in poly I: C mice, CBD 30 mg/kg reduced the activity of c-Fos+ neurons in the PFC, increased the amplitude of miniature inhibitory postsynaptic currents (mIPSCs), and reduced excitatory currents (mEPSCs) in pyramidal neurons, effectively shifting the E/I balance towards inhibition. These effects were mediated by blockade of the GPR55 receptor in PFC pyramidal neurons, reinforcing the central role of this receptor in CBD’s actions on cortical circuits relevant to ASD.

Poleg et al. (2021) demonstrated, using magnetic resonance spectroscopy (MRS) in cerebrospinal fluid (CSF), that CBD (via Avidekel) reduced glutamate levels in the CSF of Shank3 mice without altering GABA levels. In contrast, pure THC reduced glutamate and increased GABA, suggesting complementary mechanisms of action for the two phytocannabinoids on excitatory and inhibitory neurotransmitter systems. Additionally, CBD modulated mRNA expression in the cerebellum of these animals. However, these effects require further investigation.

#### 3.6.3 Neuroinflammation and Oxidative Stress

CBD treatment reduced markers of neuroinflammation and oxidative stress in several ASD models.

Faiza Ali et al. (2025) demonstrated that CBD 100 mg/kg (chronic) increased antioxidant enzymes (GSH, SOD, catalase) and reduced markers of oxidative stress (lipid peroxidation, nitric oxide) in the hippocampus and prefrontal cortex of VPA BALB/c mice. Additionally, CBD reduced hippocampal atrophy, pyramidal neuron degeneration in the PFC, and Purkinje cell degeneration in the cerebellum, demonstrating a comprehensive neuroprotective effect.

Costa et al. (2024) also observed that CBD 100 mg/kg (chronic) reduced hippocampal oxidative stress (thiol marker) in VPA Wistar rats, without alterations in CAT and SOD enzymes or in the posterior cortex. Zamberletti et al. (2019) observed reductions in inflammatory markers (GFAP, CD11b, TNF-α) and restoration of microglial morphology in the hippocampus with CBDV at 20 mg/kg, suggesting that VPA-induced hippocampal neuroinflammation is a target responsive to cannabinoid treatment. These findings support the anti-inflammatory and neuroprotective properties of CBD.

#### 3.6.4 Transcriptomic and Molecular Modulation

Emerging evidence suggests that CBD may influence gene expression patterns associated with ASD.

Transcriptomic analyses demonstrated modulation of differentially expressed genes involved in synaptic function, neurodevelopment, immune signaling, and neuronal communication. Loomis et al. (2025) performed a CPF transcriptomic analysis following treatment with JZP541 (botanical CBD) and identified modulation of differentially expressed genes (DEGs) associated with ASD, providing the most direct evidence to date that CBD can correct specific molecular signatures of ASD at the transcriptional level. Although still limited, these findings suggest that CBD may exert effects at multiple levels of biological organization.

### 3.7 Risk of Bias Assessment

The risk-of-bias analysis revealed important methodological limitations among the included studies. The risk of bias assessment of the 14 included studies was conducted using the SYRCLE Risk of Bias Tool for animal studies (Hooijmans et al., 2014), encompassing ten domains organized into six categories: selection bias (sequence generation, baseline characteristics, and allocation concealment), performance bias (random allocation and experimenter blinding), detection bias (randomized outcome assessment and evaluator blinding), attrition bias (incomplete outcome data), reporting bias (selective outcome reporting), and other sources of bias. The risk-of-bias plots were generated using the Robvis tool (McGuinness & Higgins, 2021) and are presented as unweighted bar plots because no quantitative meta-analysis was performed in this review (Figures 2 and 3).

**Figure 2.**
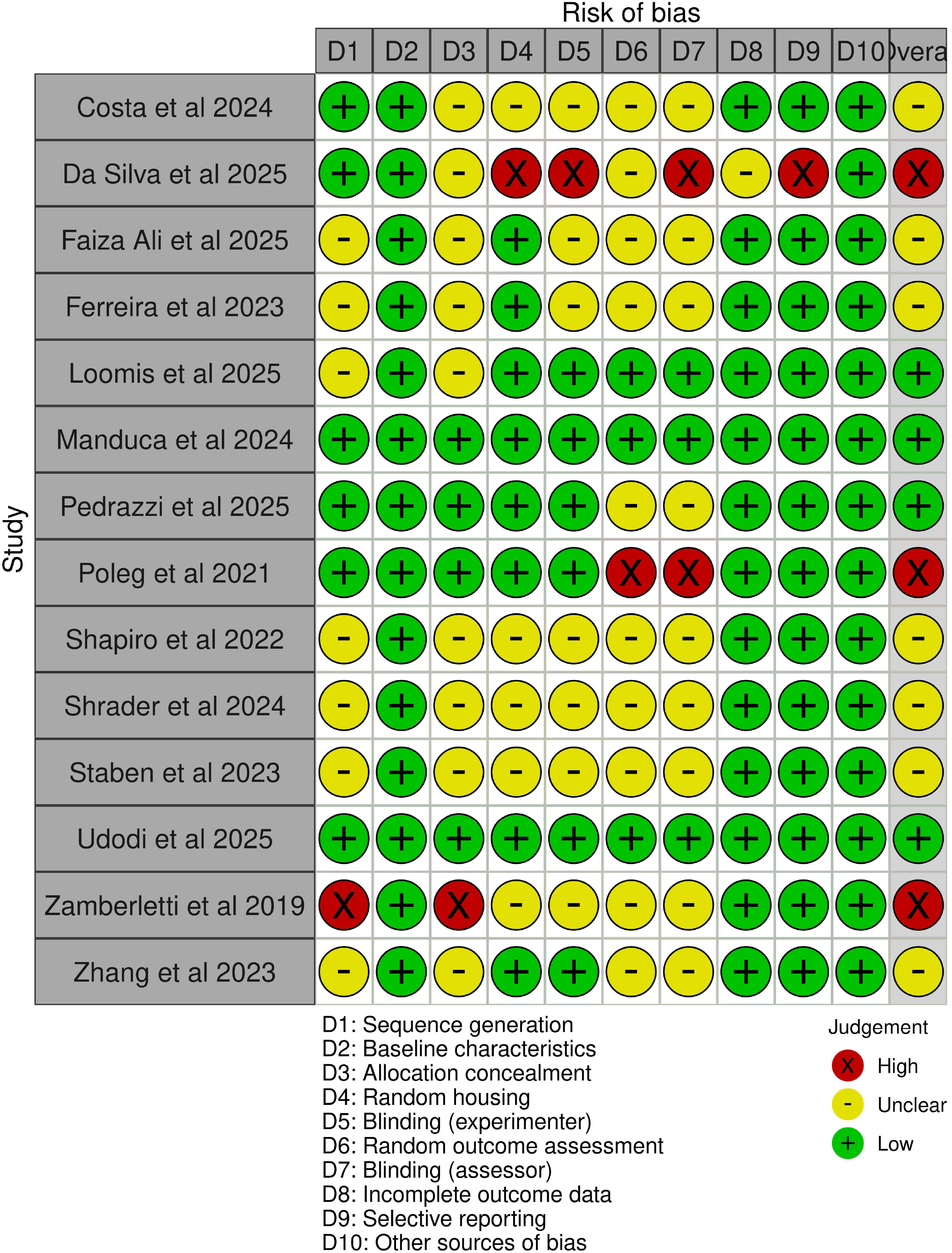
Detailed risk-of-bias assessment for individual studies using the SYRCLE Risk of Bias Tool. Green (+): low risk of bias; Yellow (-): unclear risk of bias; Red (X): high risk of bias.

**Figure 3.**
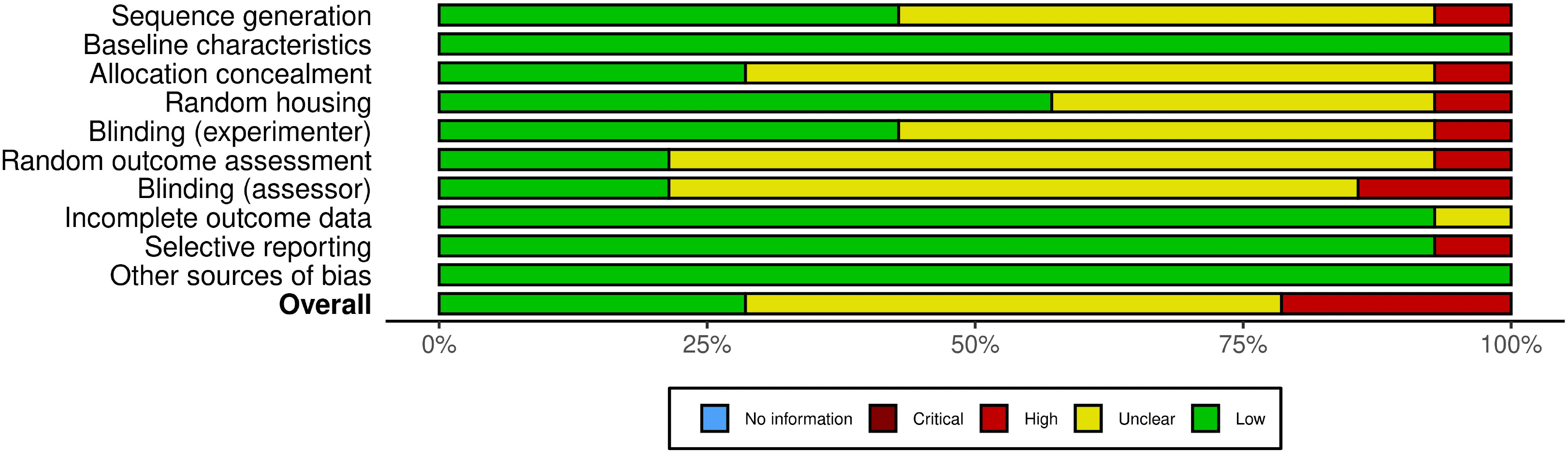
Distribution of risk of bias by domain according to the SYRCLE Risk of Bias Tool. Bars represent the percentage of studies classified as low risk (green), unclear risk (yellow), high risk (red), critical risk (dark red), and no information (blue).

In the sequence generation domain, six studies were classified as having a low risk of bias (Da Silva et al., 2025; Pedrazzi et al., 2025; Udodi et al., 2025; Costa et al., 2024; Manduca et al., 2024; Poleg et al., 2021), seven presented an uncertain risk (Faiza Ali et al., 2025; Loomis et al., 2025; Shrader et al., 2024; Ferreira et al., 2023; Staben et al., 2023; Zhang et al., 2023; Shapiro et al., 2022), and one was classified as having a high risk of bias (Zamberletti et al., 2019). In the baseline characteristics domain, all 14 studies were classified as having a low risk of bias, indicating that the experimental groups were adequately comparable at the beginning of the experiments. In the allocation concealment domain, four studies presented low risk (Pedrazzi et al., 2025; Udodi et al., 2025; Manduca et al., 2024; Poleg et al., 2021), nine were classified as uncertain risk (Da Silva et al., 2025; Faiza Ali et al., 2025; Loomis et al., 2025; Costa et al., 2024; Shrader et al., 2024; Ferreira et al., 2023; Staben et al., 2023; Zhang et al., 2023; Shapiro et al., 2022), and one presented high risk (Zamberletti et al., 2019).

In the random allocation domain, eight studies were classified as having a low risk of bias (Faiza Ali et al., 2025; Loomis et al., 2025; Pedrazzi et al., 2025; Udodi et al., 2025; Manduca et al., 2024; Ferreira et al., 2023; Zhang et al., 2023; Poleg et al., 2021), five presented an uncertain risk (Costa et al., 2024; Shrader et al., 2024; Staben et al., 2023; Shapiro et al., 2022; Zamberletti et al., 2019), and one was classified as having a high risk (Da Silva et al., 2025). In the experimenter blinding domain, six studies presented low risk (Loomis et al., 2025; Pedrazzi et al., 2025; Udodi et al., 2025; Manduca et al., 2024; Zhang et al., 2023; Poleg et al., 2021), seven were classified as uncertain risk (Faiza Ali et al., 2025; Costa et al., 2024; Shrader et al., 2024; Ferreira et al., 2023; Staben et al., 2023;; Shapiro et al., 2022; Zamberletti et al., 2019), and one presented high risk (Da Silva et al., 2025).

In the randomized outcome assessment domain, three studies were classified as low risk (Loomis et al., 2025; Udodi et al., 2025; Manduca et al., 2024), ten presented uncertain risk (Da Silva et al., 2025; Faiza Ali et al., 2025; Pedrazzi et al., 2025; Costa et al., 2024; Shrader et al., 2024; Ferreira et al., 2023; Staben et al., 2023; Zhang et al., 2023; Shapiro et al., 2022; Zamberletti et al., 2019), and one was classified as high risk (Poleg et al., 2021). In the domain of evaluator blinding, three studies presented low risk (Loomis et al., 2025; Udodi et al., 2025; Manduca et al., 2024), nine were classified as uncertain risk (Faiza Ali et al., 2025; Pedrazzi et al., 2025; Costa et al., 2024; Shrader et al., 2024; Ferreira et al., 2023; Staben et al., 2023; Zhang et al., 2023; Shapiro et al., 2022; Zamberletti et al., 2019) and two presented high risk (Da Silva et al., 2025; Poleg et al., 2021).

In the domain of incomplete outcome data, 13 studies were classified as having a low risk of bias, with only Da Silva et al. (2025) presenting an uncertain risk, and no studies classified as high risk.

Similarly, in the domain of selective outcome reporting, 13 studies were classified as low risk, with Da Silva et al. (2025) the only study with a high risk of bias. In the domain of other sources of bias, all 14 studies were classified as low risk, indicating no additional relevant sources of bias not covered by the other domains.

In general, the risk of bias varied among the included studies. Udodi et al. (2025) and Manduca et al. (2024) were the studies with the best methodological profile, presenting a low risk of bias in all domains evaluated. Da Silva et al. (2025) and Poleg et al. (2021) presented the most domains at high risk of bias.

Regarding selection, performance, and detection bias, not all studies reported whether animals were randomly assigned to experimental groups or whether researchers responsible for the care and application of treatments were aware of each animal’s intervention group, potentially influencing the conduct of the experiment. This is an important limitation, considering behavioral tests such as the Marble Burying and grooming test depend on a researcher’s direct observation. On the positive side, all 14 studies demonstrated that the groups were similar at the beginning of the experiment, an important aspect for the validity of the results.

Attrition and reporting bias-wise, however, had satisfactory results regarding data completeness and reporting. Most studies presented the data in full, without unjustified exclusions, and no evidence was found that unfavorable results were omitted.

No study presented other relevant sources of bias either. Overall, these findings indicate that, despite limitations related to blinding and randomization, the reported data appear to be complete and transparent and should be considered when interpreting the results of this review. The risk-of-bias assessment indicates the need for greater methodological rigor and transparency in future preclinical studies evaluating cannabinoid-based interventions in ASD models.

## 4. Discussion

### 4.1 Behavioral and Neurobiological Outcomes

The present systematic review aimed to synthesize and analyze the methodological quality of the available evidence regarding the psychopharmacological effects of CBD and CBD-related formulations in rodent animal models of ASD. Evidence from 14 preclinical studies suggests that CBD exerts beneficial effects across multiple behavioral domains relevant to ASD, particularly social interaction, repetitive behaviors, anxiety-like behaviors, and cognitive performance. Furthermore, several studies identified neurobiological effects, including modulation of the endocannabinoid system, restoration of excitation/inhibition (E/I) balance, anti-inflammatory and antioxidant actions, and transcriptional regulation, despite methodological heterogeneity among studies.

Among all behavioral domains investigated, social behavior emerged as the most consistently improved outcome following CBD administration. Improvements were observed across pharmacological models induced by prenatal exposure to valproic acid (VPA), genetic models such as BTBR, Scn8a, Shank3, and Fmr1, and maternal immune activation models induced by poly I: C. Since social deficits represent one of the core diagnostic features of ASD, these findings are particularly relevant from a translational perspective. Symptoms such as anxiety, stereotypies, and social and cognitive deficits in animal models with ASD are related to imbalances between arousal and inhibition (E/I) and dysfunctions in the endocannabinoid system, particularly in CB1 and CB2 receptors (Zamberletti, Gabaglio, & Parolaro, 2017). The observation that CBD improved social interaction across distinct etiological models suggests that its therapeutic actions may target convergent neurobiological pathways involved in social dysfunction rather than model-specific mechanisms.

Repetitive and stereotyped behaviors were also consistently attenuated by CBD. Reductions in grooming, marble burying, hyperlocomotion, and stereotyped activity were reported in multiple ASD models. These findings are of particular interest because repetitive behaviors are thought to involve dysfunction within cortico-striatal-thalamic circuits, which are strongly influenced by endocannabinoid signaling. In this context, there is evidence that modulation of the endocannabinoid system can directly influence the exacerbated behaviors present in ASD. According to Campanale, Siniscalco, and Di Marzo (2025), inhibition of the enzymes FAAH (Fatty Acid Amide Hydrolase) and MAGL (Monoacylglycerol Lipase), responsible for the degradation of the main endocannabinoids anandamide (AEA) and 2-arachidonoylglycerol (2-AG), respectively, can improve repetitive, stereotyped behaviors and sensory alterations, as well as reduce deficits in sociability, anxiety, and depressive symptoms in experimental models of ASD. The authors also highlight the involvement of the endocannabinoid system in synaptic plasticity and neuroinflammation, mechanisms already implicated in the pathophysiology of ASD.

Anxiety-like behaviors constitute one of the most prevalent psychiatric comorbidities associated with ASD (Mcvey, 2019). A recent systematic review indicates that the prevalence of anxiety in individuals with ASD can range from 0% to 82.2%, being one of the most frequently reported psychiatric comorbidities in this population (Bougeard et al., 2021). In this scenario, several studies reporting anxiolytic effects of CBD in the Elevated Plus Maze, Open Field, and Marble Burying paradigms are relevant findings. Interestingly, some studies reported dose-dependent responses, with intermediate doses yielding greater behavioral improvements than lower or higher doses.

This pattern is consistent with the inverted U-shaped dose-response curve frequently described for CBD in both preclinical and clinical research, not restricted to ASD but also observed in studies of chronic pain, comorbid depression, and panic (Brito et al., 2025; Malvestio et al., 2021; De Freitas et al., 2010). These findings together reinforce that the absence of effect at certain doses should not be interpreted as ineffectiveness of the compound, but as evidence of the need for careful titration and rigorous definition of therapeutic windows for each condition investigated.

The neurobiological findings synthesized in this review suggest that CBD may act through multiple complementary mechanisms. Only seven of the 14 studies investigated the neurobiological mechanisms of CBD, a relatively low proportion. They indicated four main CBD routes of action: modulation of the endocannabinoid system, restoration of excitation/inhibition (E/I) balance, anti-inflammatory and antioxidant activity, and transcriptional modulation.

Several studies reported modulation of the endocannabinoid system, including changes in CB1, CB2, FAAH, MAGL, and GPR55 signaling pathways (Udodi et al., 2025; Manduca et al., 2024; Zamberletti et al., 2019). Given the central role of the endocannabinoid system in synaptic plasticity, emotional regulation, social behavior, and neurodevelopment, these findings provide strong biological plausibility for the therapeutic effects of CBD in ASD.

Another important observation was the convergence of evidence supporting restoration of excitation/inhibition (E/I) balance. Alterations in E/I homeostasis are currently considered among the most influential neurobiological hypotheses for ASD pathophysiology. Zhang et al. (2023) demonstrated that CBD reduced glutamatergic activity and increased GABAergic activity in prefrontal cortex neurons of poly I: C-treated mice, thereby favoring greater neuronal balance. Similarly, Udodi et al. (2025) observed increased expression of parvalbumin-positive (PV+) GABAergic interneurons after CBD treatment in VPA mice. This finding is relevant because the reduction of PV+ neurons has already been described in both individuals with ASD and animal models of the condition, and is associated with the E/I imbalance characteristic of the disorder (Filice et al., 2020). The restoration of intraneuronal parvalbumin-expressing populations and the modulation of GPR55-dependent pathways further support this mechanism. The role of this receptor in emotional and cognitive functions was explored in Marichal-Cancino et al.’s (2017) review, which suggests decreased GPR55 expression, particularly in regions such as the hippocampus and frontal cortex, in animals with ASD. However, the data requires further investigation.

Additionally, several studies identified anti-inflammatory and antioxidant effects associated with CBD treatment. Neuroinflammation and oxidative stress have emerged as relevant targets, since prenatal exposure to VPA is known to induce alterations in brain systems that contribute to the development of the autistic phenotype in offspring (Schneider & Przewłocki, 2005). Reductions in GFAP, CD11b, TNF-α, oxidative stress markers, and neuronal degeneration were observed in multiple ASD models, and the ability of CBD to attenuate these processes may represent an important mechanism underlying its therapeutic actions. Studies have reported reductions in hippocampal neuroinflammatory markers and restoration of microglial morphology in Zamberletti et al. (2019), increases in antioxidant enzymes and reductions in oxidative stress in Faiza Ali et al. (2025), and reductions in hippocampal oxidative stress in Costa et al. (2024).

Emerging evidence also suggests that CBD may influence transcriptional regulation. The transcriptomic findings reported by Loomis et al. (2025) demonstrated modulation of differentially expressed genes associated with ASD following treatment with a CBD-rich botanical formulation. Although still limited, these results suggest that CBD may act at multiple levels of biological organization, ranging from neurotransmission to gene expression.

It is important to note that the neurobiological mechanisms reported by the 14 included studies do not fully encompass all of CBD’s pharmacological effects. Traditionally, CBD acts as an agonist of the 5-HT_1A_ serotonergic receptor, a mechanism widely associated with its anxiolytic and antidepressant effects (Acunha et al., 2026), and as an activator of the TRPV_1_ channel, which is involved in calcium modulation (Costa et al., 2004). In this review, none of the included studies specifically addressed these pathways in the context of ASD, representing a significant gap in the preclinical literature, given the characteristics of the autistic phenotype and the implications of the 5-HT_1A_ and TRPV_1_ pathways.

### 4.2 Pharmacological Considerations

A key secondary finding in this review was the compositional diversity of CBD, with favorable effects observed not only in isolated preparations but also across various formulations and in combination with other chemical components. Since effective exposure to CBD depends on the route of administration, the vehicle, and the formulation technology (Moazen-Zadeh et al., 2024; Grifoni et al., 2022), this diversity is relevant for the interpretation of outcomes.

Recent reviews on this topic, such as that by Vasconcelos et al. (2026), have focused on the behavioral pharmacology of isolated CBD in animal models of ASD. Although the primary objective of the present review was also to evaluate the effect of isolated CBD, the search identified studies that used CBD in different formulations, compositions, and combinations with other drugs. These studies met the eligibility criteria and were included; the diversity of interventions was treated as a variable in interpreting the outcomes, rather than as a methodological limitation. This secondary finding broadens our understanding of the potential uses of CBD in rodent models of ASD and justifies the analysis conducted.

#### 4.2.1 Pharmacological Diversity as an Experimental Variable

Among the included studies, the interventions encompassed pure CBD; CBD-rich botanical extracts such as JZP541; mixed formulations containing THC (Avidekel); terpene blends; structural analogs such as cannabidivarin (CBDV); a nanostructured formulation (CBDON); and combination therapy with an antipsychotic. The effects of these compounds should be interpreted with caution and on a case-by-case basis, as they do not represent variations within the same therapeutic category but rather interventions with distinct pharmacological targets.

THC acts as a direct partial agonist of the CB1 and CB2 receptors (Pertwee, 2008), directly activating the CB1 receptor and contributing to specific behavioral effects (anxiogenic or sedative) that cannot be attributed to CBD alone. As discussed earlier in the Introduction section of this review, CBD has very low affinity for CB1 and acts as a negative allosteric modulator of this receptor (Laprairie et al., 2015; Thomas et al., 2007). Thus, extracts such as Avidekel, which contain THC, even at low concentrations, can produce behavioral effects resulting from a pharmacological interaction or combination. Terpenes can also exert effects of their own and modify cannabinoid activity (LaVigne et al., 2021). In the literature, this phenomenon is described as the “entourage effect” (Russo, 2011). Although they are of therapeutic interest, these combined effects make it impossible to isolate each component’s contribution to the outcomes and limit comparisons with isolated CBD.

CBDV, in turn, is an analog of CBD, structurally identical except for the length of the alkyl side chain (Hill et al., 2013). Like CBD, CBDV also has low affinity for the CB1 receptor; however, it acts predominantly on TRPV1 and TRPA1 channels, the GPR55 receptor, and enzymes involved in endocannabinoid metabolism (Cascio; Pertwee 2014). Many of these targets are shared with CBD, but the profile of potency, efficacy, and selectivity may differ.

Furthermore, nanoemulsions have been developed to increase the solubility, stability, and drug release, typically at low doses (20 to 200 nanometers), thereby accelerating the absorption of these compounds (Preeti et al., 2023; Moazen-Zadeh et al., 2024; Grifoni et al., 2022). For CBD, a lipophilic molecule with low aqueous solubility and extensive first-pass metabolism, this technology directly addresses its pharmacokinetic limitations and drives the development of strategies like this one. The nanoemulsion represents a promising alternative, but the data do not permit a conclusion regarding its superiority over the isolated compound, as only one study in this review included this technology.

The combination of CBD and risperidone resulted in improved sociability, an effect not observed with CBD alone at the same dose, suggesting that CBD may act as an adjunct rather than solely as monotherapy. This finding has significant translational relevance, as the pharmacological management of symptoms associated with ASD relies on antipsychotics, the use of which is limited by adverse effects; an adjunctive role for CBD could, in theory, allow for a reduction in the dose of these medications, a possibility not tested in the included studies.

#### 4.2.2 Routes of Administration, Bioavailability, and Dose Variability

Different routes of administration were mentioned in the studies included in this review, including intraperitoneal, oral gavage, and inhalation. In humans, the oral and intraperitoneal routes undergo first-pass hepatic metabolism, resulting in notably low bioavailability (Millar et al., 2018). This pharmacokinetic characteristic is also present with gavage and intraperitoneal administration in rodents, meaning that the effects of exposure may be less than the chosen dose would actually suggest. This may explain, to some extent, the choice of high doses in the experimental studies (up to 100 mg/kg) aimed at circumventing first-pass metabolism and achieving significant behavioral effects. However, the included studies do not state this rationale. On the other hand, the inhalation route differs from this process, favoring higher plasma concentrations and possibly greater penetration of the blood-brain barrier; however, exposure via passive inhalation is difficult to quantify.

This review also revealed a wide range of doses (0.1-100 mg/kg), which may partly reflect the pharmacokinetic differences among the routes of administration and formulations examined here. The absence of standardized protocols in preclinical research on TEA makes direct comparisons between studies difficult. Differences in species, strain, and treatment duration may further contribute to the variability observed across studies. Furthermore, the loss of effect in some cases, both above and below a certain dose range, indicates nonlinear dose-response relationships. This implies that the effective dose window is specific to each preparation and that the efficacy of a formulation must be evaluated relative to its own control with its own dose-response curve, rather than inferred from the doses of the isolated compound.

Taken together, the results indicate that CBD interventions in rodent models of ASD do not constitute a homogeneous category and that composition, formulation, route, dose, and administration all influence exposure and the pharmacological profile. The potential benefits of alternatives to the isolated compound, such as greater efficacy per dose unit, synergistic effects, and contributions from other constituents, are, for now, hypotheses consistent with the data and cannot be considered advantages in and of themselves. Future research should systematically evaluate pharmacokinetic parameters and dose-response relationships by comparing isolated CBD, these alternatives, and their respective controls, using the same route of administration, model, and set of outcomes, to establish therapeutic regimens that are relevant from a translational perspective.

### 4.3 Limitations and Methodological Quality

Several critical factors must be considered when interpreting the findings of this review. Foremost, as previously noted, substantial heterogeneity was observed across the included studies concerning the animal models utilized, CBD formulations, treatment protocols, dosages, routes of administration, treatment duration, and targeted behavioral outcomes. This high degree of methodological variability precluded a quantitative synthesis via meta-analysis.

Second, the risk-of-bias assessment conducted using the SYRCLE tool revealed widespread reporting deficiencies in randomization procedures, allocation concealment, and blinding across several studies. The domains of experimenter and evaluator blinding, and the random assignment of animals to the tests, presented the highest proportion of uncertain risk. Only the studies by Udodi et al. (2025) and Manduca et al. (2024) showed low risk in all domains. The absence of blinding is especially relevant in behavioral tests dependent on direct observation, as such limitations can compromise study findings and lead to overestimation of effect sizes (Voelkl et al., 2018; Landis, 2012).

Future studies should prioritize rigorous methodological standardization, particularly the harmonization of behavioral protocols, dosing regimens, and outcome reporting frameworks. Further investigation is also warranted to elucidate sex-dependent differences and under-explored neurobiological pathways, such as 5-HT_1A_ and TRPV_1_ signaling, in the context of ASD and CBD. Addressing these gaps will be key to determining whether CBD and related compounds can move beyond symptomatic management to serve as disease-modifying strategies for ASD.

## 5. Conclusion

The available preclinical evidence suggests that CBD-based interventions exert beneficial psychopharmacological effects across diverse rodent models of ASD. In the majority of included studies, these interventions improved social behavior, reduced repetitive behaviors, attenuated anxiety-like symptoms, and, in certain instances, enhanced cognitive performance. These behavioral improvements were underpinned by neurobiological changes, including modulation of the endocannabinoid system, restoration of the excitation/inhibition balance, anti-inflammatory and antioxidant actions, neuroprotection, and transcriptional regulation.

Although this review primarily focused on isolated CBD, several included studies also evaluated CBD in various alternative formulations, including plant extracts, structural analogs, and combinations with terpenes or synthetic antipsychotics. While these findings are promising, the outcomes observed with multicomponent preparations cannot be attributed solely to CBD. Furthermore, the purported advantages of these complex formulations and combinations over isolated CBD have yet to be conclusively demonstrated; for instance, the nanoemulsion and the antipsychotic combination were each assessed in only a single study, without direct comparisons with isolated CBD.

Collectively, the synthesized evidence highlights cannabidiol as a highly promising candidate for further investigation, demonstrating the capacity to modulate the behavioral, molecular, and neurophysiological alterations associated with ASD. Nevertheless, significant methodological limitations persist across the literature. These include profound heterogeneity in experimental protocols, CBD compositions, formulations, dosages, and administration routes, alongside insufficient risk-of-bias reporting and a pervasive underrepresentation of female subjects.

While CBD emerges as a compelling therapeutic strategy for ASD, rigorously designed preclinical and clinical investigations are imperative to fully establish its efficacy, delineate its mechanisms of action, determine optimal dosing regimens, and confirm its translational relevance. Future preclinical research must systematically evaluate pharmacokinetic parameters and dose-response relationships by directly comparing isolated CBD with its complex formulations and combinations, alongside appropriate controls, within the same experimental framework. Ultimately, integrating behavioral, electrophysiological, molecular, and transcriptomic approaches will be essential to ascertain whether the promising effects observed in rodent models can be successfully translated into effective clinical interventions for individuals with ASD.

### Strengths and Limitations of the Review

The present review possesses several notable strengths. To the best of our knowledge, it is the first systematic review specifically designed to evaluate the psychopharmacological effects of cannabidiol in rodent models of autism spectrum disorder (ASD) while simultaneously examining underlying neurobiological mechanisms and methodological quality. Furthermore, the review strictly adhered to PRISMA 2020 guidelines, employed a comprehensive search strategy across multiple databases (including gray literature), and systematically assessed risk of bias using the SYRCLE tool.

Nevertheless, certain limitations must be acknowledged. The included studies exhibited profound heterogeneity regarding ASD models, cannabinoid formulations, dosages, treatment durations, routes of administration, and behavioral assays. Consequently, a quantitative synthesis via meta-analysis was unfeasible. Finally, pervasive methodological limitations were identified across multiple studies, most notably the inadequate reporting of randomization and blinding procedures.

## # DECLARATIONS

## ## Acknowledgements

The authors thank the University of São Paulo (USP), the Ribeirão Preto Medical School (FMRP-USP), the Ribeirão Preto School of Philosophy, Sciences and Letters (FFCLRP-USP), the Federal University of Paraíba (UFPB), and the Federal University of Alfenas (UNIFAL-MG) for their institutional support.

The authors also acknowledge all researchers whose studies have advanced the understanding of cannabidiol and autism spectrum disorder.

## ## Author Contributions

Conceptualization: Bruna Rabelo Ribeiro Domingues, Renato Leonardo de Freitas.

Methodology: Bruna Rabelo Ribeiro Domingues, Gabriel Santos Félix, Maria Fernanda Lucena Bezerra, Renato Leonardo de Freitas.

Literature Search and Study Selection: Bruna Rabelo Ribeiro Domingues, Gabriel Santos Félix, Maria Fernanda Lucena Bezerra.

Data Extraction and Risk of Bias Assessment: Bruna Rabelo Ribeiro Domingues, Gabriel Santos Félix, Maria Fernanda Lucena Bezerra.

Data Analysis and Interpretation: Bruna Rabelo Ribeiro Domingues, Priscila Medeiros, Renato Leonardo de Freitas.

Writing – Original Draft Preparation: Bruna Rabelo Ribeiro Domingues.

Writing – Review and Editing: Bruna Rabelo Ribeiro Domingues, Priscila

Medeiros, Renato Leonardo de Freitas.

Supervision: Renato Leonardo de Freitas.

All authors read and approved the final manuscript.

## ## Data Availability Statement

All data generated or analyzed during this study are included in this published article and its supplementary information files.

## ## Ethics Approval

Ethical approval was not required because this study is a systematic review based exclusively on previously published literature and did not involve direct experimentation on humans or animals.

## ## Consent for Publication

Not applicable.

## Supplementary material - Search strategy

Search terms were combined using the Boolean operators “OR” and “AND”. The search strategy presented below was applied across the PubMed, Web of Science (Clarivate Analytics), Embase, and Scopus databases, as well as grey literature sources, including ProQuest Dissertations & Theses.

